# UCM-A86 is a selective positive allosteric modulator of GluN1/GluN3 NMDA receptors

**DOI:** 10.1101/2025.11.14.688378

**Authors:** Avery J. Benton, Mia R. Johns, Emil Diamant, Nirvan Rouzbeh, Valdemar P. Sørensen, Carly M. Anderson, Davide Secci, Zhucheng Zhang, Andrew R. Rau, Rasmus P. Clausen, Kasper B. Hansen

## Abstract

*N*-methyl-D-aspartate (NMDA) receptors are ionotropic glutamate receptors that mediate excitatory neurotransmission in the central nervous system (CNS) where they play critical roles in normal and pathological brain functions and neurodevelopment. While the glutamate/glycine-activated GluN2-containing NMDA receptors (GluN1/GluN2) have been extensively studied, the physiological roles and pharmacology of glycine-activated GluN3-containing receptors (GluN1/GluN3) remain less understood. Although GluN1/GluN3 receptors exhibit unique functional properties and play distinct roles in neuronal development and synapse maturation, studies of their precise roles in neurophysiology and circuit function are impeded by limited availability of GluN3-selective pharmacological tools. This study describes UCM-A86, a novel GluN3-selective positive allosteric modulator, with EC_50_ values of 21 µM and 19 µM at GluN1/GluN3A and GluN1/GluN3B receptors, respectively. UCM-A86 selectively potentiates recombinant GluN1/GluN3A and GluN1/GluN3B receptors by 436% and 174%, respectively, relative to activation by glycine, with no activity at recombinant GluN1/GluN2A-D receptors. Furthermore, UCM-A86 selectively potentiates responses from native GluN1/GluN3A receptors expressed in somatostatin-expressing interneurons of the somatosensory cortex with no modulation of hippocampal AMPA receptor- and GluN1/2 NMDA receptor-mediated excitatory postsynaptic currents. Mechanistic studies suggest that UCM-A86 modulation is facilitated by agonist binding (or channel gating) and that UCM-A86 primarily potentiates GluN1/GluN3A by increasing open probability with no effects on mean channel conductance. These findings advance the synthetic pharmacology of GluN1/GluN3 receptors and provide a novel tool for modulation of native GluN3-containing NMDA receptors.

**Significance statement:** This study introduces UCM-A86 as the first positive allosteric modulator selective for GluN3-containing NMDA receptors, addressing a critical gap in the pharmacological toolbox for investigating these understudied receptor subtypes. Using electrophysiological approaches in both recombinant and native systems, UCM-A86 demonstrates specific modulation of GluN3-containing NMDA receptors without affecting GluN2-containing NMDA receptors or AMPA receptors. UCM-A86 therefore provides new avenues to investigate the physiological roles of GluN3 subunits in normal and pathological brain function.

## Introduction

N-methyl-D-aspartate (NMDA) receptors are ligand-gated cation channels that play crucial roles in excitatory neurotransmission throughout the central nervous system (CNS) (Hansen et al., 2021). Neuronal NMDA receptors are fundamental to numerous neurobiological processes including neurodevelopment and synaptic plasticity, and inappropriate distribution, regulation, or biophysical function of NMDA receptors contribute to the pathophysiology of multiple neurological and psychiatric disorders (Hansen et al., 2021; Nicoll and Schulman, 2023; Beaurain et al., 2024; Hanson et al., 2024).

There are seven NMDA receptor subunits, namely GluN1, GluN2A-D, and GluN3A-B, but the predominant heterotetrameric NMDA receptor complex in the brain is composed of two glycine/D-serine-binding GluN1 and two glutamate-binding GluN2 subunits (Hansen et al., 2018; Hansen et al., 2021). These GluN1/GluN2 receptors have been extensively studied, leading to the development of numerous modulators with therapeutic potential as treatments in several CNS diseases (Hansen et al., 2021; Hanson et al., 2024). However, neuronal NMDA receptors can also assemble as GluN1/GluN3 receptors, comprising two GluN1 and two glycine/D-serine-binding GluN3 subunits (Crawley et al., 2022; Michalski and Furukawa, 2024). While GluN1/GluN2 receptors show high calcium permeability, pronounced magnesium block, and require both glycine and glutamate binding for activation (Hansen et al., 2018; Hansen et al., 2021), GluN1/GluN3 receptors display low calcium permeability, minimal magnesium block, and can be activated by glycine alone (Chatterton et al., 2002; Crawley et al., 2022). Additionally, GluN1/GluN3 receptors exhibit strong desensitization in response to glycine, in contrast to the relatively weak desensitization observed in GluN1/GluN2 receptors following activation by glutamate and glycine (Awobuluyi et al., 2007; Kvist et al., 2013; Grand et al., 2018; Rouzbeh et al., 2023).

The GluN3A subunit plays important roles in synapse maturation and plasticity, with high and widespread expression early in development (Chatterton et al., 2002; Wong et al., 2002; Crawley et al., 2022; Gonzalez-Gonzalez et al., 2023). Although the widespread expression of GluN3A declines into adolescence and adulthood, expression levels remain elevated in specific cell types, including somatostatin-expressing interneurons (Wong et al., 2002; Perez-Otano et al., 2016; Bossi et al., 2022; Crawley et al., 2022; Bossi et al., 2023). Deletion of GluN3A increases hippocampal spine density and disrupts activity-dependent synapse stabilization, particularly during early developmental periods (Perez-Otano et al., 2006; Marco et al., 2013; Yuan and Bellone, 2013; Kehoe et al., 2014; Perez-Otano et al., 2016). Dysregulation of GluN3A has been implicated in various neurological and psychiatric disorders, including Huntington’s disease (Marco et al., 2013; Mahfooz et al., 2016), schizophrenia (Matsuno et al., 2015; Greenwood et al., 2016), addiction (Yuan et al., 2013; Christian et al., 2021), and fear- and anxiety-related behaviors (Conde-Dusman et al., 2021; Bossi et al., 2022; Pizzamiglio et al., 2025), highlighting the importance of developing novel pharmacological modulators targeting GluN1/GluN3A receptors for potential therapeutic applications. In contrast to GluN3A, much less is known about the physiological roles of the GluN3B subunit that is primarily expressed in motor neurons of the brainstem and spinal cord during development and into adulthood (Chatterton et al., 2002; Matsuda et al., 2003; Fukaya et al., 2005).

Despite the importance of GluN1/GluN3A in neuronal functions, the synthetic pharmacology of GluN3-containing receptors is underdeveloped compared to the pharmacological toolbox available for subunit-selective allosteric modulation of GluN2-containing NMDA receptors. To date, EU1180-438 and WZB117 have been described as negative allosteric modulators with selectivity for GluN1/GluN3 over GluN1/GluN2 receptors (Zhu et al., 2020; Zeng et al., 2022). The unique activation properties of GluN1/GluN3 receptors, in which glycine binding to GluN3 results in receptor activation and glycine binding to GluN1 facilitates receptor desensitization, endow GluN1-selective competitive antagonists with the potential to act as positive allosteric modulators by reducing receptor desensitization (Madry et al., 2007; Grand et al., 2018; Rouzbeh et al., 2023). According to this mechanism, CGP 78608 and L-689,560 potentiate glycine-activated responses from GluN1/GluN3 receptors but also strongly inhibit GluN1/GluN2 receptors (Grand et al., 2018; Rouzbeh et al., 2023). Despite these recent advances, positive allosteric modulators with selectivity for GluN1/GluN3 over GluN1/GluN2 receptors have not been described.

Here, we uncover the mechanism of action and pharmacological activity of UCM-A86 as a novel GluN3-selective positive allosteric modulator. Using two-electrode voltage-clamp electrophysiology, we show that UCM-A86 selectively potentiates responses from GluN1/GluN3A/B over GluN1/GluN2A-D receptors. Mouse brain slice electrophysiology corroborates this selectivity by demonstrating potentiation of native GluN1/GluN3A NMDA receptors in somatostatin-expressing interneurons without affecting synaptic responses from hippocampal AMPA or GluN1/2 NMDA receptors. Fast-application whole-cell patch-clamp electrophysiology experiments provide insights into the mechanism of action for UCM-A86 that highlights the complex structure-function relationship of GluN1/GluN3 receptors.

## Materials and Methods

### DNA constructs and ligands

Plasmids with cDNAs encoding rat GluN1-1a (Genbank accession numbers U11418 and U08261), GluN1-1b (U08263), GluN1-4a (U08267), GluN1-4b (U08268), GluN2A (D13211), GluN2B (U11419), GluN2C (M91563), GluN2D (L31611), GluN3A (U29873), and GluN3B (AF440691) were used for NMDA receptor expression in *Xenopus laevis* oocytes and HEK293T cells. For expression in *Xenopus* oocytes, the cDNA was linearized using restriction enzymes and used for *in vitro* cRNA synthesis (mMessage mMachine, Thermo Fisher Scientific, Waltham, MA; HiScribe T7 ARCA mRNA kit, New England Biolabs, Ipswich, MA). To enable cRNA synthesis by the T7 RNA polymerase for expression in *Xenopus* oocytes, the rat GluN2B cDNA was modified to remove a T7 RNA polymerase termination site located in the C-terminal domain as previously described (Hansen et al., 2014). In some experiments with GluN1/GluN3 receptors, GluN1 subunits included the mutations in the agonist binding site, F484A + T518L in GluN1-a splice variants (GluN1-1a^FATL^ or GluN1-4a^FATL^) and F505A + T539L in GluN1-b splice variants (GluN1-1b^FATL^ or GluN1-4b^FATL^), to prevent glycine binding and desensitization (Kvist et al., 2013). UCM-A86 was identified in a virtual screening campaign as previously described (Atomwise, 2024) and subsequently purchased from Enamine (Monmouth Junction, NJ) or synthesized (See Supplemental Information) to be used in experiments with >95% purity. All other ligands were obtained from Sigma-Aldrich (St. Louis, MO), Tocris Bioscience (Minneapolis, MN), or Hello Bio (Princeton, NJ). The stock solution for UCM-A86 was made at 20 mM, 30 mM, or 100 mM in DMSO and the stock solution for CGP 78608 was made at 10 mM in water with 22 mM NaOH. DMSO concentrations were kept constant in recording solutions and never exceeded 0.5%.

### Animals

All experimental procedures using animals were approved by the University of Montana Institutional Animal Care and Use Committee and were performed in accordance with state and federal Animal Welfare Acts and the policies of the Public Health Service. Adult mice aged 6-8 weeks of both sexes were used for experiments. All mice were group housed on a 12:12 light/dark cycle and were given access to food and water at libitum. Wild-type (WT) (C57Bl/6J; strain 000664) and mice expressing enhanced green fluorescent protein (EGFP) in SST interneurons (Oliva et al., 2000) (hereafter referred to as GIN; strain 003718) mice were purchased from The Jackson Laboratory (Bar Harbor, ME). GIN mice were backcrossed into the C57Bl/6J genetic background for >9 generations.

### Two-electrode voltage-clamp electrophysiology

NMDA receptors were expressed in *Xenopus* oocytes obtained from Xenopus 1 Corporation (Dexter, MI) by injecting cRNAs encoding GluN1 and GluN2 or GluN3 subunits essentially as previously described (Rouzbeh et al., 2023). Oocytes were incubated for 2-5 days at 17°C before two-electrode voltage-clamp recordings were performed at room temperature (21–23°C) as previously described (Hansen et al., 2013). During recordings, oocytes were perfused with extracellular recording solution containing (in mM) 90 NaCl, 1 KCl, 10 HEPES, 0.01 EDTA, 0.5 BaCl_2_ (pH 7.4 with NaOH). In addition, 2-hydroxypropyl-β-cyclodextrin (5 mM) was included in the extracellular recording solution to enhance solubility and minimize potential issues related to compound precipitation. Precipitation of UCM-A86 due to limited solubility was not observed at concentrations up to 300 µM (higher concentrations were not evaluated). Recording electrodes were filled with 3.0 M KCl and current responses were recorded at a holding potential of −40 mV. Responses were measured using a two-electrode voltage-clamp amplifier (OC725C; Warner Instruments, Holliston, MA), low-pass filtered at 20 Hz (Alligator Technologies, Charlottesville, VA), and digitized using a PCI-6025E data acquisition board (National Instruments, Austin, TX). Solutions were applied by gravity-driven perfusion through digital 8-modular valve positioners (Hamilton Company, Reno NV).

### Whole-cell patch-clamp electrophysiology

Whole-cell patch-clamp electrophysiology was performed using HEK293T cells that had been transfected ∼24 hours prior to recordings using the calcium phosphate precipitation method (Chen and Okayama, 1987) with a 1:1 ratio of plasmid cDNAs encoding GluN1-1a and GluN3A. EGFP for visualization of transfected cells was expressed as a separate protein from the same plasmid as GluN1 as previously described (Yi et al., 2018). Cells were maintained in culture medium composed of Dulbecco’s modified Eagle’s medium (DMEM) with GlutaMax-I and sodium pyruvate (Thermo Fisher Scientific) supplemented with 10% fetal bovine serum (Neuromics, Edina, MN), 10 U/ml penicillin, and 10 mg/ml streptomycin (Thermo Fisher Scientific). On the day of experiments, cells were detached using Hank’s balanced salt solution (Life Technologies Coorperations) and DMEM >3 hours before recordings, and then re-plated onto glass coverslips coated with poly-D-lysine (0.1 mg/ ml) and maintained in culture medium. Recordings were performed at room temperature (21–23°C) at a holding potential of −60 mV using an Axopatch 200B amplifier (Molecular Devices, Sunnyvale, CA). Recordings were filtered at 8 kHz (8-pole Bessel, Frequency Devices, Ottawa, IL) and digitized at 20 kHz using Digidata 1322B with pClamp 10 software (Molecular Devices). Recording electrodes with tip resistances of ∼2-3 MΩ were pulled from thin-walled borosilicate glass micropipettes (Sutter Instruments, Novato, CA) using a horizontal micropipette puller (P-1000; Sutter Instruments), fire polished, and filled with internal solution containing (in mM) 110 D-gluconate, 110 CsOH, 30 CsCl, 5 HEPES, 4 NaCl, 0.5 CaCl2, 2 MgCl_2_, 5 BAPTA, 2 NaATP, and 0.3 NaGTP (pH 7.35 with CsOH). The extracellular recording solution was comprised of (in mM) 10 HEPES, 150 NaCl, 3 KCl, and 0.5 CaCl_2_ (pH to 7.4 with NaOH). Unless otherwise stated, the holding potentials were not corrected for the liquid junction potential, which was previously measured to be +10.1 ± 0.6 mV (n = 4) (Yi et al., 2018). Upon entering whole-cell recording configuration, cells were lifted off the poly-D-lysine coverslips and into the gravity-driven perfusion solution. Rapid solution exchange around the lifted cells was performed with a four-barrel, three-barrel, or theta-glass pipette controlled by a piezoelectric translator (MXPZT-300; Siskiyou Corporation, Grants Pass, OR). The open-tip solution exchange times (10-90% rise time) were typically measured to be 0.3–0.8 ms. For measurements of mean channel conductance using nonstationary variance analysis, the responses were activated by bath-application of 10 mM glycine in the absence (vehicle, 0.15% DMSO) or presence of 30 µM UCM-A86, but in the continuous presence of 1 µM CGP 78608. For all recordings, only cells with current response amplitudes of less than 1000 pA and access resistance less than 10 MΩ were used for data analyses.

### Brain slice electrophysiology

Following the induction of a deep anesthetic plane with isoflurane, WT and GIN mice (6-8 weeks) were cardiac perfused with ice-cold high sucrose solution containing (in mM) 3 KCl, 24 NaHCO_3_, 1.25 NaH_2_PO_4_, 10 glucose, 230 sucrose, 0.5 CaCl_2_, and 10 MgSO_4_. Brains were removed and coronal slices (300 µm) containing the somatosensory cortex or dorsal hippocampus were cut on a Leica VT 1200S vibratome in a solution containing (in mM) 130 NaCl, 3 KCl, 24 NaHCO_4_, 1.25 NaH_2_PO_4_, 10 glucose, 1 CaCl_2_, and 3 MgSO_4_. Brain slices were incubated in this solution for >1 hour prior to recordings, and all solutions were saturated with 95% O_2_/5% CO_2_.

Brain slices containing the hippocampus or somatosensory cortex were transferred to a recording chamber (Warner Instruments) mounted on a SliceScope Pro 2000 microscope (Scientifica, Uckfield, United Kingdom). Slices were continuously perfused with an oxygenated aCSF recording solution containing (in mM) 130 NaCl, 3 KCl, 24 NaHCO_3_, 1.25 NaH_2_PO_4_, 10 glucose, 2 CaCl_2_, and 1 MgSO_4_ at a flow rate of 2-3 ml/min. The recording solution was maintained at 32°C with a dual in-line heater/platform (Warner Instruments). Recording electrodes were pulled with a horizontal pipette puller (P-1000, Sutter Instruments) to obtain resistances of 2-3 MΩ. All recordings were made using a Multiclamp 700B amplifier (Molecular Devices), filtered at 4 kHz, and digitized at 20 kHz using Digidata 1440A with pClamp 10 software (Molecular Devices). Access resistance was monitored throughout all recordings via a −5 mV hyperpolarizing voltage jump (100 ms), and only neurons with a series resistance < 20 MΩ that did not change by more than 20% were used for data analysis.

For measurements of native GluN1/GluN3 receptor responses, somatostatin-expressing interneurons in the somatosensory cortex of GIN mice (6-8 weeks) were targeted using recording electrodes filled with internal solution containing (in mM) 120 Cs-methanesulfonate, 4.6 MgCl_2_, 10 HEPES, 15 BAPTA, 4 Na_2_-ATP, 0.4 Na-GTP, 1 QX-314, and 10 K_2_-creatine phosphate (pH 7.25 with CsOH; 280-290 mOsm). The somatostatin-expressing interneurons were visually identified by their expression of EGFP (Oliva et al., 2000). To pharmacologically isolate GluN1/GluN3-mediated current responses, 10 mM glycine was pressure applied (100 ms duration, 5 psi) using a custom-made picospritzer (openspritzer; (Forman et al., 2017) in the presence of 10 µM gabazine, 2 µM NBQX, 100 µM DL-APV, 50 µM strychnine to block GABA_A_, AMPA, GluN1/2 NMDA, and glycine receptors, respectively. After gaining access to the cell, >5 min of baseline recordings were collected while pressure-applying glycine at an interstimulus interval of 30 s. Next, 1 µM CGP 78608 was bath-applied to prevent glycine-mediated desensitization of GluN1/GluN3A receptors (Grand et al., 2018; Rouzbeh et al., 2023). After stable peak amplitudes of glycine-mediated responses were established in the presence of CGP 78608, either 20 µM UCM-A86 or vehicle (0.1% DMSO) were bath-applied for >10 min.

To assess the selectivity of UCM-A86, whole-cell responses from CA1 pyramidal neurons in acute brain slices from adult WT mice (6-8 weeks) were recorded in the presence of 2 µM NBQX and 10 µM gabazine. Recording electrodes were filled with solution containing (in mM) 120 Cs-methanesulfonate, 15 CsCl, 10 tetraethylammonium chloride, 10 HEPES, 8 NaCl, 3 Mg-ATP, 1.5 MgCl_2_, 1 QX-314, 0.3 Na-GTP, and 0.2 EGTA (pH 7.3 with CsOH); 295-305 mOsm). Pyramidal cells were clamped at a holding potential of +40 mV and NMDA receptor-mediated excitatory post synaptic currents (NMDAR-EPSCs) were evoked via delivery of an electrical stimulus (0.2 ms) along the Schaffer collaterals using a bipolar stimulating electrode at an interstimulus interval of 30 sec. The stimulus intensity was adjusted to obtain peak amplitudes of NMDAR-EPSCs at approximately 50% of the maximal response. Following a baseline period of 5-10 min, 20 µM UCM-A86 was bath-applied for 10-15 min, and 3-5 consecutive sweeps from end of this time period were averaged for data analysis. AMPA receptor-mediated EPSCs (AMPAR-EPSCs) were collected in a similar manner except that NBQX was replaced with 100 µM DL-APV in the recording solution and pyramidal cells were voltage clamped a holding potential of −70 mV.

### Data analysis

Concentration-response data were analyzed using GraphPad Prism (GraphPad Software, La Jolla, CA). Data from each individual oocyte were fit to the Hill equation:

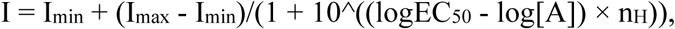

where I_min_ and I_max_ are the fitted minimum and maximum responses, respectively, to the agonist or allosteric modulator, [A] is the concentration of agonist or allosteric modulator, n_H_ is the Hill slope, and EC_50_ is the concentration of agonist or allosteric modulator that produces half-maximum response. For agonists, the fitted I_min_ is typically negligible and close to zero, and for allosteric modulators, the fitted I_min_ is typically close to amplitude of the control response to agonist measured in the absence of modulator (100%). For graphical representation, data points from individual cells were normalized and the averaged data points were plotted with the fitted concentration-response curve. The reported parameters are averages of values determined from individual oocytes (i.e., each oocyte represents one replicate).

Macroscopic responses from whole-cell patch-clamp and brain slice recordings were analyzed using Clampfit (Molecular Devices) or Axograph X (Axograph Scientific, Sydney, Australia). Rise times were measured as the duration from 10% to 90% of the maximal response amplitude. The deactivation time course was determined using two-exponential fits to obtain τ_fast_, τ_slow_, and % fast is the fitted percentage of the fast component. Weighted time constants were calculated as:

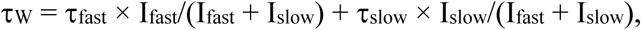

where I_slow_ and I_fast_ are fitted response amplitudes for the slow and fast components, respectively.

Nonstationary variance analysis was performed on the deactivation phase of current responses from recombinant receptors activated by glycine in the presence of 1 µM CGP 78608 and either vehicle (0.15% DMSO) or 30 µM UCM-A86 at a holding potential of −60 mV. The deactivation phase was divided into 100 equally spaced segments, and the mean current amplitude of each segment was determined. In addition, the current was high pass filtered with a cutoff of 1 Hz, and the current variance was determined for the same 100 segments. The variance as a function of mean current was fitted by the equation:

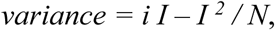

where I is the current amplitude of the response, *i* is the mean unitary current, and N is the number of channels. Mean channel conductance was calculated from the mean unitary current assuming 0 mV of reversal potential and using a holding potential that was corrected for the liquid junction potential (i.e., V_holding_ = −60 mV - (+10 mV) = −70 mV) (see Yi et al. (2018) for determination of liquid junction potential). The reported parameters are averages of values determined from individual cells (i.e., each cell represents one replicate).

### Statistical analysis

Experiments were exploratory and designed to assess biological hypotheses without a predetermined number of replicates. Experimental data were collected on at least two different days using freshly made recording solutions with multiple batches of transfected HEK293T cells for whole-cell patch-clamp recordings and multiple batches of oocytes for two-electrode voltage-clamp recordings. Unless otherwise stated, the data are presented at mean ± SEM and n is the sample size that represent independent biological replicates (i.e., the number of cells). EC_50_ values are presented with their 95% confidence interval (CI) derived from the corresponding logEC_50_ values. Statistical comparisons of time constants (i.e., τ values) were performed on normally distributed rate constants (1/τ) and statistical comparisons of EC_50_ values were performed on normally distributed logEC_50_ values (Christopoulos, 1998). Significance was set at p < 0.05 and statistical analysis was performed using GraphPad Prism (GraphPad Software, La Jolla, CA) as described in figure and table legends.

## Results

### UCM-A86 is a positive allosteric modulator of GluN1/GluN3 receptors

To evaluate the mechanism of action for UCM-A86, we measured responses activated by UCM-A86 (30 µM) alone, UCM-A86 (30 µM) + glycine (30 µM), and glycine (30 µM) alone at GluN1-1a^FATL^/3A receptors using two-electrode voltage-clamp electrophysiology (Fig. 1A-C). GluN1-1a^FATL^/3A includes two mutations in the GluN1 agonist binding pockets (F484A + T518L) that abolish glycine binding and stabilize the GluN1 agonist binding domain in a conformation that prevent receptor desensitization (Kvist et al., 2013). UCM-A86 alone did not activate GluN1-1a^FATL^/3A but potentiated responses to 313 ± 22% (n = 7) relative to responses activated by glycine (100 %) (Fig. 1B,C). These results demonstrate that UCM-A86 is not an allosteric or orthosteric agonist, but rather increases GluN1/GluN3A receptor currents as a positive allosteric modulator.

**Figure 1.**
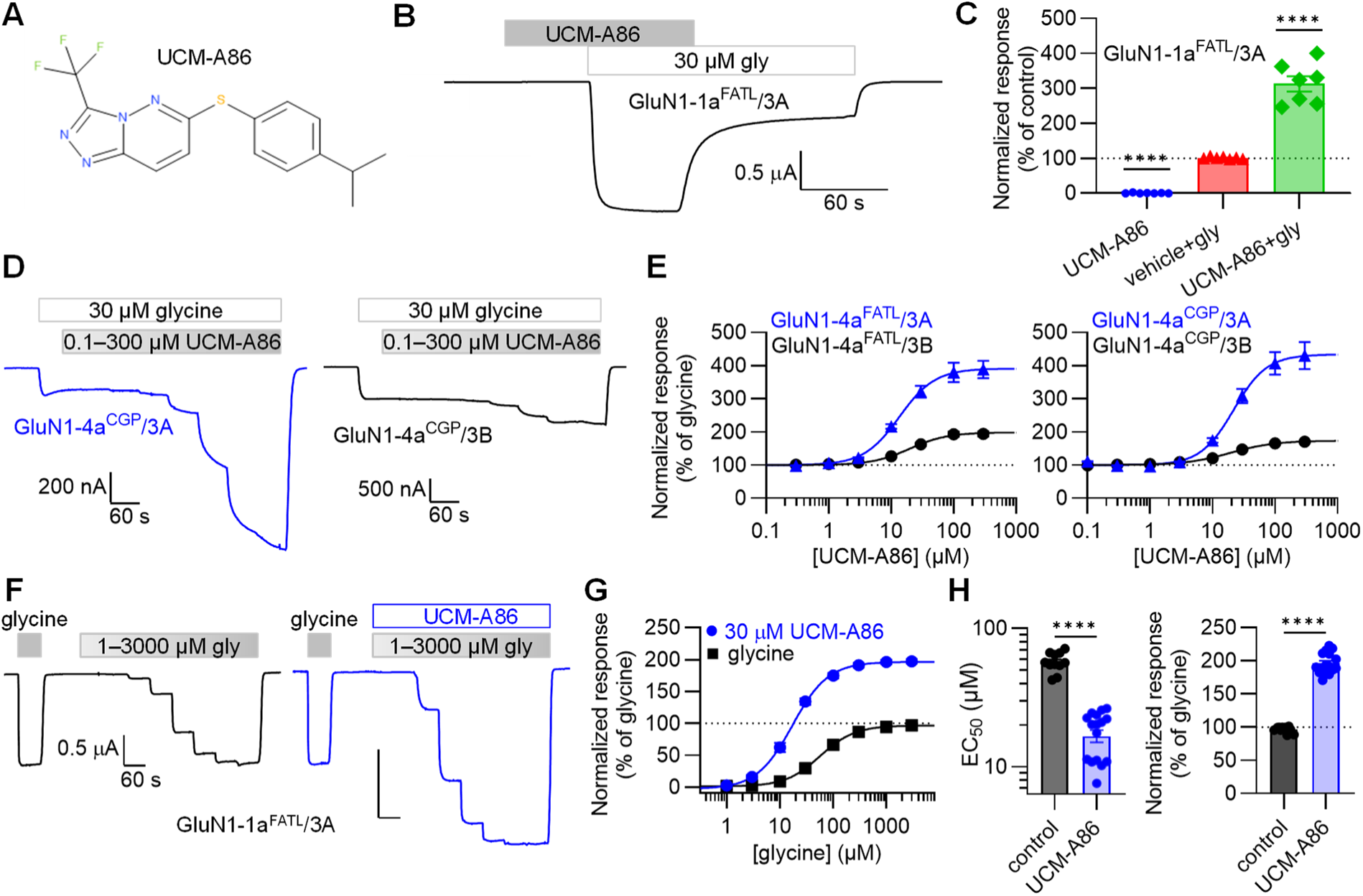
Effects of UCM-A86 on recombinant GluN3A-containing NMDA receptors. (A) Chemical structure of UCM-A86. (B) Representative two-electrode voltage-clamp recording of modulation by 30 µM UCM-A86 at recombinant GluN1-1a^FATL^/3A receptors. (C) The bar graph summarizes the responses shown in B. Responses were normalized to the glycine only response (100%). Data are mean ± SEM from 7 oocytes. Significance determined by one-sample t-test compared to 100% (****, p < 0.0001). (D) Representative recordings of UCM-A86 concentration-responses data at GluN1-4a^CGP^/3A and GluN1-4a^CGP^/3B receptors. (E) GluN1-4a^FATL^/3A-B and wildtype GluN1-4a^CGP^/3A-B receptors were activated by glycine (30 µM) in the absence and presence of increasing concentrations of UCM-A86. Data are mean ± SEM from 10 to 16 oocytes. (F) Representative recordings evaluating glycine concentration-response data in the presence and absence of 30 µM UCM-A86. (G) Glycine concentration-response data ± 30 µM UCM-A86 for GluN1-1a^FATL^/3A receptors normalized to initial response to 1 mM glycine (100% control) in the absence of UCM-A86. Data are mean ± SEM from 12 to 16 oocytes. (H) Bar graphs show the maximum responses and EC_50_ values in the absence (control) and presence of UCM-A86. Statistical significance was analyzed using unpaired t-test (****, p < 0.0001). See Table 1 for maximum responses and EC_50_ values.

**Table 1.** UCM-A86 modulation of GluN1/GluN3 receptor subtypes.

|  | GluN3A |  |  |  | GluN3B |  |  |  |
| --- | --- | --- | --- | --- | --- | --- | --- | --- |
|  | EC<sub>50</sub> (μM)<br>(95% CI) | Hill<br>slope | %<br>potentiation | n | EC<sub>50</sub> (μM)<br>(95% CI) | Hill<br>slope | %<br>potentiation | n |
| GluN1-1a<sup>FATL</sup> | 17<br>(15 - 18) | 1.6 | 396 ± 25 * | 12 | - |  |  |  |
| GluN1-1b<sup>FATL</sup> | 13 *<br>(10 - 17) | 1.6 | 367 ± 25 * | 13 | - |  |  |  |
| GluN1-4a<sup>FATL</sup> | 12 *,#<br>(9 - 15) | 1.7 | 392 ± 29 # | 16 | 20 #<br>(18 - 23) | 1.6 | 195 ± 8 # | 11 |
| GluN1-1a<sup>CGP</sup> | 24<br>(20 - 28) | 1.7 | 558 ± 34 * | 13 | - |  |  |  |
| GluN1-1b<sup>CGP</sup> | 21 *<br>(17 - 25) | 1.6 | 511 ± 48 * | 10 | - |  |  |  |
| GluN1-4a<sup>CGP</sup> | 21 *<br>(19 - 24) | 1.7 | 436 ± 41 # | 11 | 19<br>(16 - 23) | 1.3 | 174 ± 3 # | 10 |

To determine the potency and efficacy of UCM-A86, we measured concentration-response data from GluN1-4a^FATL^/3A and GluN1-4a^FATL^/3B receptors activated by 30 µM glycine (Fig. 1D). UCM-A86 potentiated responses from GluN1-4a^FATL^/3A with an EC_50_ of 12 µM (95% CI, 9-15 µM) and a maximal potentiation of 392 ± 29% (n = 16) and GluN1-4a^FATL^/3B with an EC_50_ of 20 µM (95% CI, 18-23 µM) and a maximal potentiation of 195 ± 8% (n = 11) (Fig. 1D-E and Table 1). Furthermore, we characterized UCM-A86 at wildtype GluN1/GluN3 receptors in the continuous presence of the GluN1-selective competitive antagonist CGP-78608 (hereafter CGP) that minimize desensitization by preventing glycine binding to GluN1 (Grand et al., 2018; Rouzbeh et al., 2023). UCM-A86 potentiated responses from GluN1-4a^CGP^/3A with an EC_50_ of 21 µM (95% CI, 19-24 µM) and a maximal potentiation of 436 ± 41% (n = 11) and GluN1-4a^CGP^/3B with an EC_50_ of 19 µM (95% CI, 16-23 µM) and a maximal potentiation of 174 ± 3% (n = 10) (Fig. 1D-E and Table 1). Although significant differences were observed between EC_50_ values at GluN1^FATL^- versus GluN1^CGP^-containing receptors and at GluN3A- versus GluN3B-containing receptors, the differences were minor (Table 1). Thus, UCM-A86 did not discriminate between GluN1/GluN3A and GluN1/GluN3B in terms of potency, but displayed higher efficacy at GluN1/GluN3A compared to GluN1/GluN3B. The similar activity of UCM-A86 at GluN1^FATL^/GluN3 versus GluN1^CGP^/GluN3 receptors demonstrated that allosteric modulation is not mediated by binding to the mutated GluN1 agonist binding pocket (i.e., F484A + T518L) as previously demonstrated for L-689,560 (Rouzbeh et al., 2023).

We measured glycine concentration-response data in the absence and presence of UCM-A86 at GluN1-1a^FATL^/3A receptors using two-electrode voltage-clamp electrophysiology (Fig. 1F-G). Glycine EC_50_ significantly decreased from 57 µM (95% CI, 51-63 µM; n = 11) in vehicle (0.15% DMSO) to 17 µM (95% CI, 13-21 µM; n = 16) in presence of 30 µM UCM-A86 (Fig. 1H). Under these conditions, the maximal potentiation by UCM-A86 was 196 ± 4% (n = 16) relative to the maximal glycine response measured in the same recording (Fig. 1H). Thus, positive allosteric modulation by UCM-A86 increases both the potency and the maximal response to glycine at GluN1-1a^FATL^/3A receptors.

### Modulation by UCM-A86 is not influenced by GluN1 splice variants

The GluN1 subunit exists in 8 different isoforms due to alternative splicing in the extracellular N-terminal domain (GluN1-Xa and GluN1-Xb) and the intracellular C-terminal domain (GluN1-1x, GluN1-2x, GluN1-3x, GluN1-4x) (Fig. 2A), and alternative splicing in the N-terminal domain of GluN1 can influence allosteric modulation in GluN1/2 receptors (Hansen et al., 2017; Hansen et al., 2018). We compared modulation by UCM-A86 for GluN3A co-expressed with either GluN1-1a, GluN1-1b, or GluN1-4a subunits. No significant differences in UCM-A86 EC_50_ values were observed between GluN1 splice variants, but the use of F484A + T518L or CGP resulted in small, but significant changes in EC_50_ values for some receptor subtypes (Fig. 2C and Table 1). No significant differences in maximal potentiation were observed between GluN1 splice variants (Fig. 2D and Table 1). Furthermore, the use of F484A + T518L or CGP to prevent desensitization did not influence maximum potentiation (Fig. 2D and Table 1). These findings demonstrate that UCM-A86 modulation of GluN1/GluN3A receptors is not markedly influenced by the GluN1 isoform present in the receptor complex.

**Figure 2.**
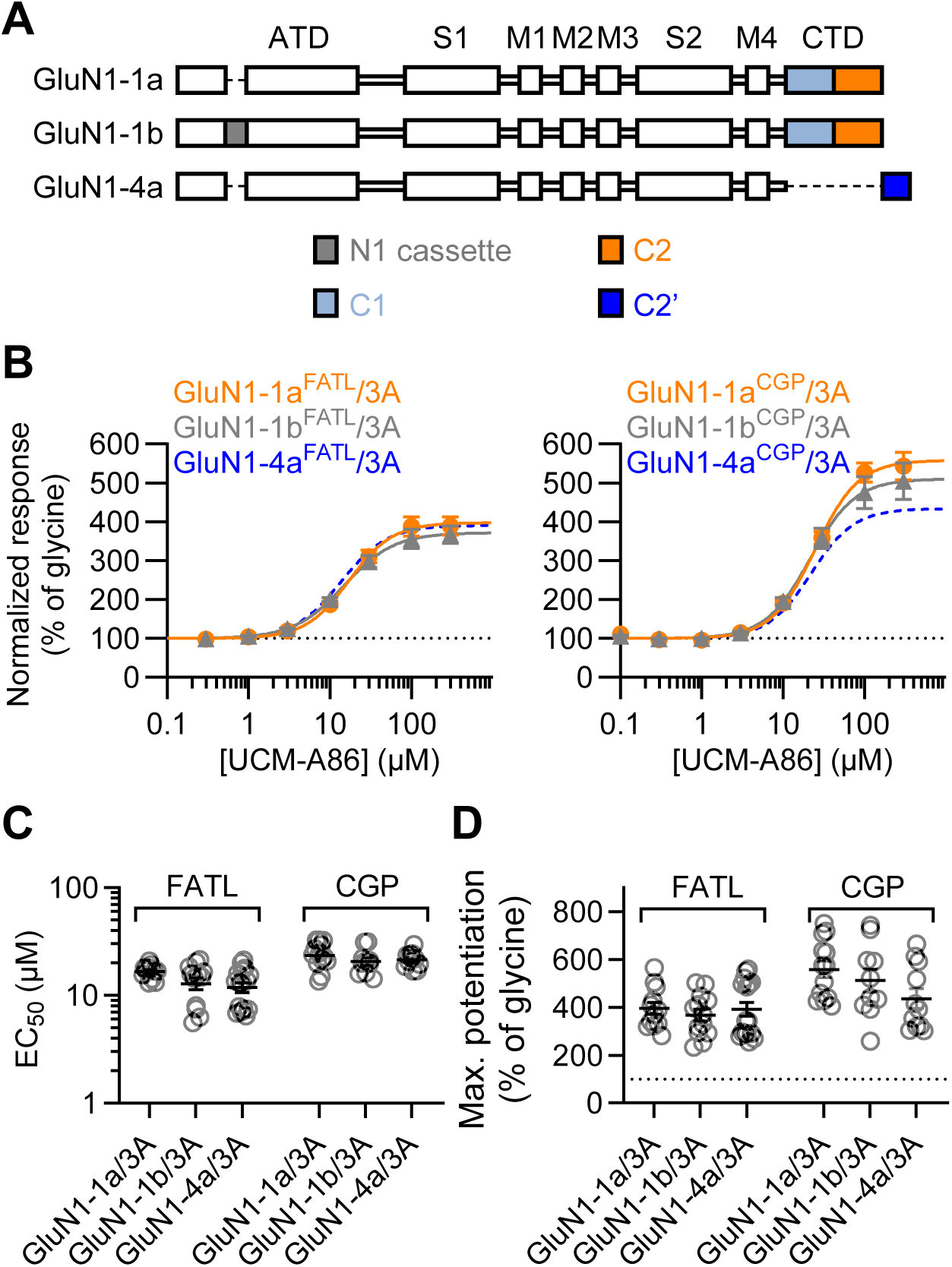
GluN1 splice variant effects on UCM-A86 modulation. (A) Schematic representation of the polypeptide chain for relevant GluN1 splice variants co-expressed with GluN3A. (B) Concentration-response data measured using two-electrode voltage-clamp electrophysiology demonstrating modulation by UCM-A86 of GluN1 splice variants co-expressed with GluN3A. F484A + T518L (FATL) mutations or CGP were used to prevent desensitization and receptors were activated by 30 µM glycine followed by increasing concentrations of UCM-A86. Data are mean ± SEM from 10 to 16 oocytes. (C-D) Summary of EC_50_ values and maximum potentiation for UCM-A86 at GluN1 splice variants co-expressed with GluN3A. See Table 1 for statistical analysis.

### UCM-A86 is selective for GluN1/GluN3 over GluN1/GluN2 receptors

To evaluate the selectivity for GluN3-containing NMDA receptors, we determined the effects of 30 µM UCM-A86 on responses from recombinant GluN1-1a/2A-D receptors activated by co-application of saturating glycine and glutamate (100 µM each) (Fig. 3). UCM-A86 slightly reduced responses to 97 ± 2% (n = 6) at GluN1-1a/2A, 92 ± 1% (n = 6) at GluN1-1a/2B, 96 ± 2% (n = 6) at GluN1-1a/2C, and 91 ± 1% (n = 6) at GluN1-1a/2D. In the same experiment, UCM-A86 potentiated GluN1^FATL^/3A receptors to 437 ± 20% (n = 8) relative to the response to 30 µM glycine alone. Thus, 30 µM UCM-A86 only produces modest, if any, modulation of recombinant GluN2-containing NMDA receptors, suggesting that UCM-A86 is selective for GluN1/GluN3 over GluN1/GluN2 receptors.

**Figure 3.**
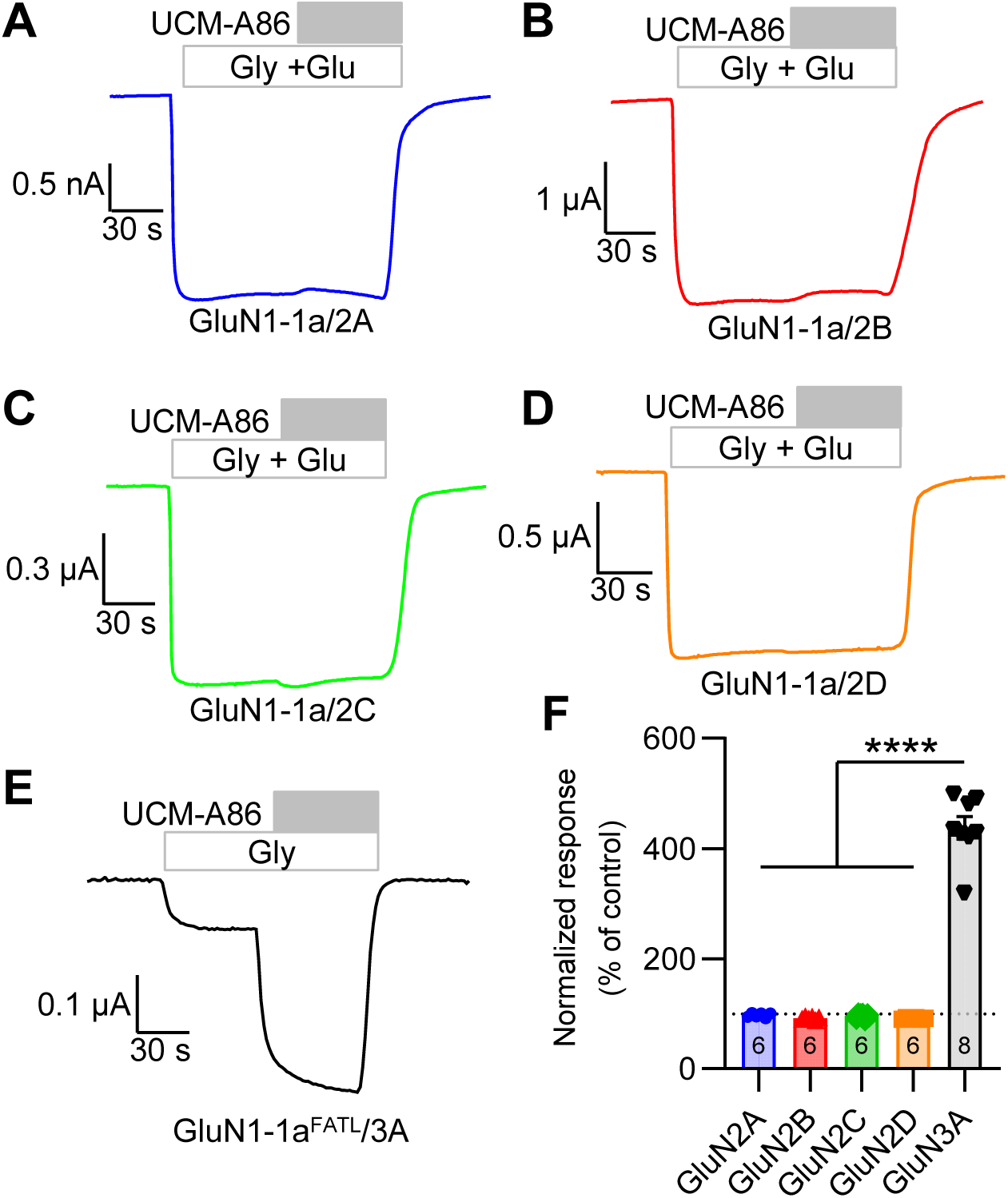
Selectivity of UCM-A86 for GluN3- over GluN2-containing NMDA receptors. (A-E) Representative two-electrode voltage-clamp recordings for GluN1-1a/2A-D receptors activated by 100 µM glycine and 100 µM glutamate (control) and GluN1-1a^FATL^/3A receptors activated by 30 µM glycine (control), followed by the addition of 30 µM UCM-A86. (F) Summary of UCM-A86 responses relative to control. Statistical significance between GluN1-1a/2A-D receptors and GluN1-1a^FATL^/3A receptors was determined using one-way ANOVA (****, p < 0.0001).

### UCM-A86 positively modulates native GluN3A-containing NMDA receptors

To evaluate the utility of UCM-A86 for modulation of native GluN3A-containing NMDA receptors, we prepared acute brain slices containing the somatosensory cortex from adult mice expressing GFP in somatostatin-expressing interneurons (Oliva et al., 2000). In the adult mouse brain, GluN3A is highly colocalized with somatostatin-expressing interneurons in multiple brain regions, including the somatosensory cortex (Bossi et al., 2022; Crawley et al., 2022; Bossi et al., 2023). Whole-cell voltage-clamp recordings were made from somatostatin-expressing interneurons visually identified based on GFP expression in the somatosensory cortex (Fig. 4A). Glycine (10 mM) was pressure-applied to the soma of recorded neurons in the presence of AMPA, GABA_A_, glycine and GluN1/2 receptor antagonists (see Materials and Methods). As in heterologous expression systems, GluN1/GluN3A receptors rapidly desensitize in the presence of glycine and responses were therefore not activated under baseline conditions (i.e. in the absence of CGP) (Fig. 4B,C). However, pressure-application of glycine resulted in GluN1/GluN3A receptor-mediated current responses when 1 µM CGP was bath-applied to the extracellular recording solution (Fig. 4B,C). Previous studies have demonstrated that these current responses are not observed in brain slices from GluN3A knockout mice or in cell types known to lack GluN3A expression (Grand et al., 2018; Zhu et al., 2020; Rouzbeh et al., 2023). Following stabilization of the glycine response, UCM-A86 (20 µM) or vehicle (0.1% DMSO) was bath-applied and UCM-A86 significantly potentiated GluN1/GluN3A receptor-mediated glycine responses (290 ± 50%; n = 8) compared to vehicle (98 ± 5%; n = 8) (relative to responses in the presence of CGP alone; unpaired t-test, p < 0.05) (Fig. 4B-E). These results demonstrate that UCM-A86 is an efficacious PAM of native GluN1/GluN3A, consistent with results from recombinant receptors using two-electrode voltage-clamp recordings.

**Figure 4.**
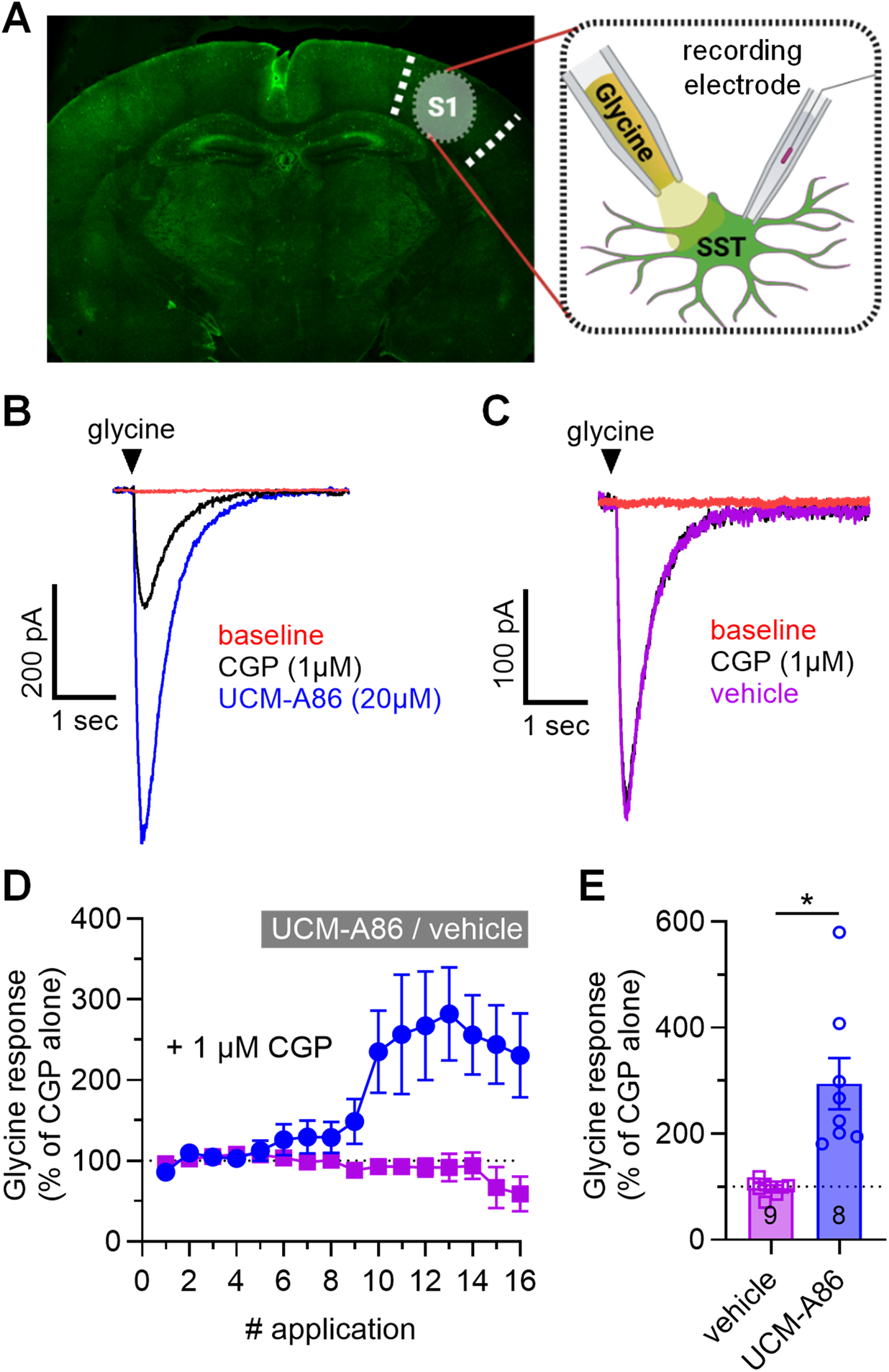
UCM-A86 potentiates glycine-mediated responses in mouse somatostatin (SST) neurons. (A) Coronal brain slice showing the primary somatosensory cortex (S1) from which whole-cell patch clamp recordings were made from GFP-expressing SST cells. (B) Representative recording demonstrating current responses from SST cells in response to pressure applied 10 mM glycine at baseline (pink), in the presence of 1 µM CGP-78608 (black), and following bath application of 20 µM AIMS 86 (blue). (C) Representative recording demonstrating current responses from SST cells in response to pressure applied 10 mM glycine at baseline (pink), in the presence of 1 µM CGP-78608 (black), and following bath application of 0.15% DMSO vehicle (purple). (D) Cumulative time course showing the effect of AIMS 86 on glycine-mediated responses. (E) Summary data demonstrates a significant potentiation of glycine-mediated currents after bath application of AIMS 86. (***, p < 0.001; unpaired t-test).

### UCM-A86 has no activity at neuronal AMPA and GluN2-containing NMDA receptors

To further corroborate the selectivity of UCM-A86 for GluN1/GluN3 receptors, we prepared acute coronal brain slices containing the dorsal hippocampus from adult mice. Whole-cell voltage-clamp recordings were made from CA1 pyramidal cells and evoked postsynaptic currents were generated by stimulation of the Shaffer collaterals (Fig. 5A). NMDA receptor-mediated EPSCs (NMDAR-EPSCs) were recorded at holding potential of +40 mV in the presence of 2 µM NBQX and 10 µM gabazine, and AMPA receptor-mediated EPSCs (AMPAR-EPSCs) were recorded at holding potential of −70 mV in the presence of 100 µM AP5 and 10 µM gabazine. Bath-application of 20 µM UCM-A86 did not significantly change the amplitude or the decay time constant of evoked NMDAR- or AMPAR-EPSCs (paired t-test, p > 0.05) (Fig. 5B-H). In summary, the results from acute brain slice preparations demonstrate that UCM-A86 selectively modulates native GluN1/GluN3A receptors with no modulatory activity on neuronal AMPA receptors or GluN1/GluN2 NMDA receptors.

**Figure 5.**
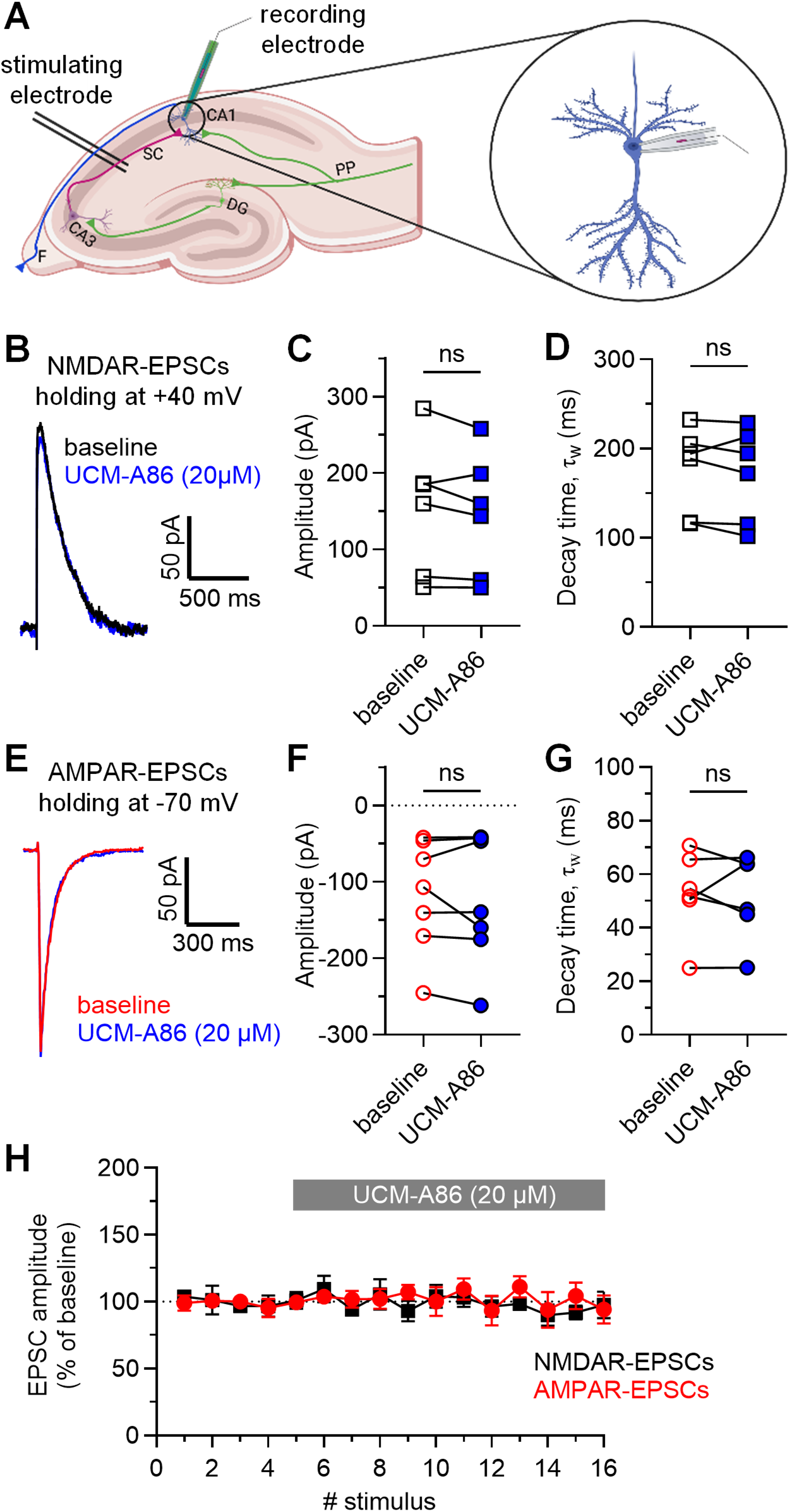
UCM-A86 lacks activity at neuronal AMPA and GluN2-containing NMDA receptors. (A) Schematic representation of the dorsal hippocampus from mouse brain. (B) Representative NMDAR-EPSCs recorded from CA1 pyramidal cells at baseline and in the presence of 20 µM UCM-A86. (C-D) UCM-A86 (20 µM) does not modulate evoked NMDAR-EPSCs in the mouse hippocampus. (ns, p>0.05; paired t-test) (E) Representative AMPAR-EPSCs recorded from CA1 pyramidal cells at baseline and in the presence of 20 µM UCM-A86. (F-G) UCM-A86 (20 µM) does not modulate evoked AMPAR-EPSCs in the mouse hippocampus. Significance determined using a paired t-test (ns, p > 0.05). (H) Cumulative time course data showing the lack of effect of 20 µM UCM- on AMPAR- and NMDAR-EPSCs.

### UCM-A86 modulation is facilitated by glycine binding or channel gating

To investigate the mechanism of action for UCM-A86, we measured allosteric modulation of GluN1-1a/GluN3A receptors expressed in HEK293T cells using fast-application whole-cell patch-clamp electrophysiology. In the continuous presence of 1 µM CGP, a long 3 s exposure of 10 mM glycine activated current responses with fast rise time (7.9 ± 0.2 ms, n = 16) to a peak amplitude (I_p_) that rapidly desensitized to a steady-state amplitude (I_ss_, 54 ± 2% of I_p_, n = 16) (Fig. 6A and Table 2). In the continuous presence of 30 µM UCM-A86, the steady-state glycine responses were potentiated to 1000 ± 250% (n = 7) relative to control with a markedly slower rise time (265 ± 77, n = 7) and a non-desensitizing response time course with no initial peak amplitude (Fig. 6A and Table 2). The mean weighted deactivation time constant (τ_w_) was 252 ± 23 ms (n = 16) for glycine responses in vehicle (0.15% DMSO) and 652 ± 74 ms (n = 7) in the continuous presence of 30 µM UCM-A86 (Fig. 6C-E and Table 2). We hypothesized that the slower rise time and absence of desensitization in the presence of UCM-A86 are resulting from a positive interaction between UCM-A86 modulation and glycine binding or channel gating. That is, modulation by UCM-A86 may not occur until after glycine binding, thereby slowing the apparent rise time and masking the initial peak amplitude. To test this hypothesis, we measured responses to long 3s applications of 10 mM glycine alone, but with pre- and post-exposures (i.e., pre/post exposure) to 30 µM UCM-A86 (Fig. 6B). These glycine responses displayed a relatively fast rise time (17 ± 2 ms, n = 9), a slow decay from the peak amplitude presumably mainly reflecting UCM-A86 unbinding, and a mean weighted deactivation time constant (τ_w_) of 586 ± 87 ms (n = 9) (Fig. 6C-E and Table 2). Furthermore, steady-state glycine responses with pre/post exposure of UCM-A86 were potentiated to 243 ± 22% (n = 9) relative to control. Thus, glycine responses with pre/post exposure of UCM-A86 display faster rise times and are potentiated less compared to responses in the continuous presence of UCM-A86, consistent with less binding of UCM-A86 in the absence of glycine binding or channel gating (Table 2). These results support the hypothesis that UCM-A86 modulation is facilitated by glycine binding or channel gating.

**Figure 6.**
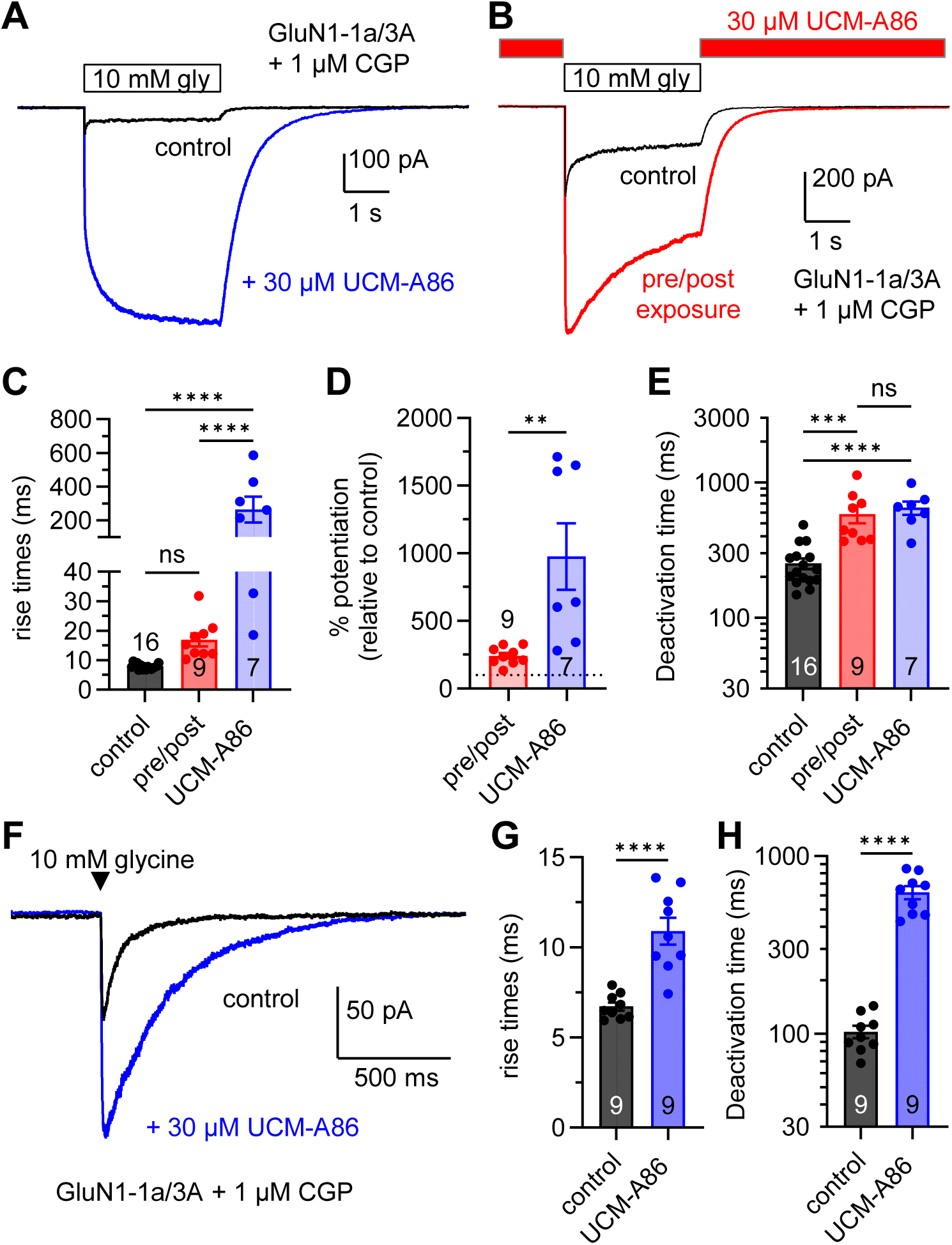
UCM-A86 modulation is facilitated by glycine binding or channel gating. (A) Representative whole-cell patch-clamp recordings demonstrating differences in the time course of responses from GluN1-1a/3A receptors activated by long (3 s) exposures to glycine and in the absence (control) or continous presence of UCM-A86. CGP-78608 (1 µM) was continously present during the recordings. (B) Representative recordings of glycine responses from GluN1-1a/3A receptors in the absence of UCM-A86 or with UCM-A86 present pre and post exposure to glycine. CGP-78608 (1 µM) was continously present during the recordings. (C) Summary of rise times for glycine responses in control, pre/post UCM-A86 exposure, and continous UCM-A86 exposure conditions. Significance determined by one-way ANOVA with Tukey post-test (****, p < 0.0001; ns, p > 0.05). (D) Summary potentiation of glycine responses in pre/post UCM-A86 exposure and continous UCM-A86 exposure conditions relative to control. Significance determined by unpaired t-test (**, p < 0.01). (E) Summary of desensitization for glycine responses in control, pre/post UCM-A86 exposure, and continous UCM-A86 exposure conditions. Significance determined by a one-way ANOVA with Tukey post-test (****, p < 0.0001; ns, p > 0.05). (F) Representative recordings of responses to brief (5 ms) exposures to glycine in the absence (control) or presence of UCM-A86. (G-H) Summaries of rise times and deactivation times for responses to brief (5 ms) glycine exposures in the absence (control) or presence of UCM-A86. Significance determined by unpaired t-test (****, p < 0.0001). See Table 2 for summary of all kinetic parameters.

**Table 2.** Macroscopic response time course for GluN1-1a/GluN3A receptors.

| Long exposure | %<br>potentiation | rise time<br>(ms) | %<br>desensitization | Deactivation time |  |  |  | n |
| --- | --- | --- | --- | --- | --- | --- | --- | --- |
| | | | | % fast | $\tau_{\text{fast}}$<br>(ms) | $\tau_{\text{slow}}$<br>(ms) | $\tau_{\text{weighted}}$<br>(ms) | |
| Glycine control | - | 7.9 ± 0.2 | 46 ± 2 | 89 ± 3 | 155 ± 9 | 1420 ± 420 | 252 ± 23 | 16 |
| UCM-A86 continuous | 1000 ± 250 | 265 ± 77 | not observed | 82 ± 6 | 433 ± 27 | 2070 ± 490 | 652 ± 74 | 7 |
| UCM-A86 pre/post | 243 ± 22 | 17 ± 2 | - | 87 ± 5 | 343 ± 29 | 3400 ± 1130 | 586 ± 87 | 9 |

| Brief exposure | %<br>potentiation | rise time<br>(ms) | %<br>desensitization | Deactivation time |  |  |  | n |
| --- | --- | --- | --- | --- | --- | --- | --- | --- |
| | | | | % fast | $\tau_{\text{fast}}$ (ms) | $\tau_{\text{slow}}$ (ms) | $\tau_{\text{weighted}}$ (ms) | |
| Glycine control | - | 6.7 ± 0.2 | - | 37 ± 5 | 56 ± 8 | 197 ± 23 | 103 ± 8 | 9 |
| UCM-A86 continuous | 292 ± 46 | 11 ± 1 | - | 26 ± 5 | 414 ± 36 | 1350 ± 110 | 626 ± 53 | 9 |

Responses from GluN1-1a/GluN3A receptors were measured using fast-application whole-cell patch-clamp recordings. Cells were activated by long (3 seconds) or short (5 ms) applications of 10 mM glycine in the continuous presence of 1 μM CGP-78608. Responses were activated in the absence (vehicle control, 0.15% DMSO) or presence of 30 μM UCM-A86. Pre/post indicates that UCM-A86 was present before and after glycine exposure, but not at the same time as glycine. Rise times are for 10-90% of the response amplitude and deactivation time constants were determined by using two-exponential fits to obtain $\tau_{\text{fast}}$ , $\tau_{\text{slow}}$ , and % fast is the fitted percentage of the fast component. Weighted time constants ( $\tau_{\text{weighted}}$ ) were calculated as described in Materials and Methods. Potentiation was measured for steady-state responses relative to glycine control during long (3s) glycine exposures and peak responses relative to glycine control during brief (5 ms) glycine exposures. Data are presented as mean ± SEM, and n is the number of cells used to generate the data. - indicates not determined.

If UCM-A86 modulation is facilitated by glycine binding, then we predict that short glycine applications would provide insufficient time for a pronounced onset of UCM-A86 modulation. To test this prediction, we measured responses to brief 5 ms applications of 10 mM glycine alone (i.e., in 0.15% DMSO vehicle) and in the continuous presence of 30 µM UCM-A86 (Fig. 6F and Table 2). Glycine alone activated peak responses with fast rise time (6.7 ± 0.2, n = 9) and mean weighted deactivation time constant (τ_w_) of 103 ± 8 ms (n = 9) (Fig. 6G,H and Table 2). Glycine in the presence of UCM-A86 activated peak responses with a slower rise time (11 ± 1, n = 9) and slower mean weighted deactivation time constant (τ_w_) of 626 ± 53 ms (n = 9). Furthermore, peak responses to brief glycine applications in the presence of UCM-A86 were potentiated to 292 ± 46% (n = 9) relative to control. Thus, UCM-A86 were unable to reach full potentiation of the peak amplitude under these conditions with brief 5 ms glycine application, consistent with our prediction and in support of the hypothesis that UCM-A86 modulation is facilitated by glycine binding or channel gating.

### UCM-A86 modulation is mediated by a multistep mechanism

To further investigate the mechanism of UCM-A86, we measured the time course for allosteric modulation of steady-state responses activated by 10 mM glycine in the continuous presence of 1 µM CGP from GluN1-1a/GluN3A receptors using fast-application whole-cell patch-clamp electrophysiology (Fig. 7A-C and Table 3). Under these conditions, both the onset and offset of modulation were best described by two-exponential fits (Table 3). Furthermore, modulation by application of either 10 µM or 30 µM UCM-A86 did not produce significant differences between the weighted time constants or the level of potentiation (Fig. 7A-C and Table 3). The presence of two components in the onset and offset of UCM-A86 modulation is different from the mono-exponential time course that would be predicted by a simple bimolecular interaction. Thus, these results suggest that UCM-A86 modulation involves a multistep and more complex mechanism that includes different binding affinities to distinct receptor states or other factors that may influence binding and efficacy at the allosteric binding site.

**Figure 7.**
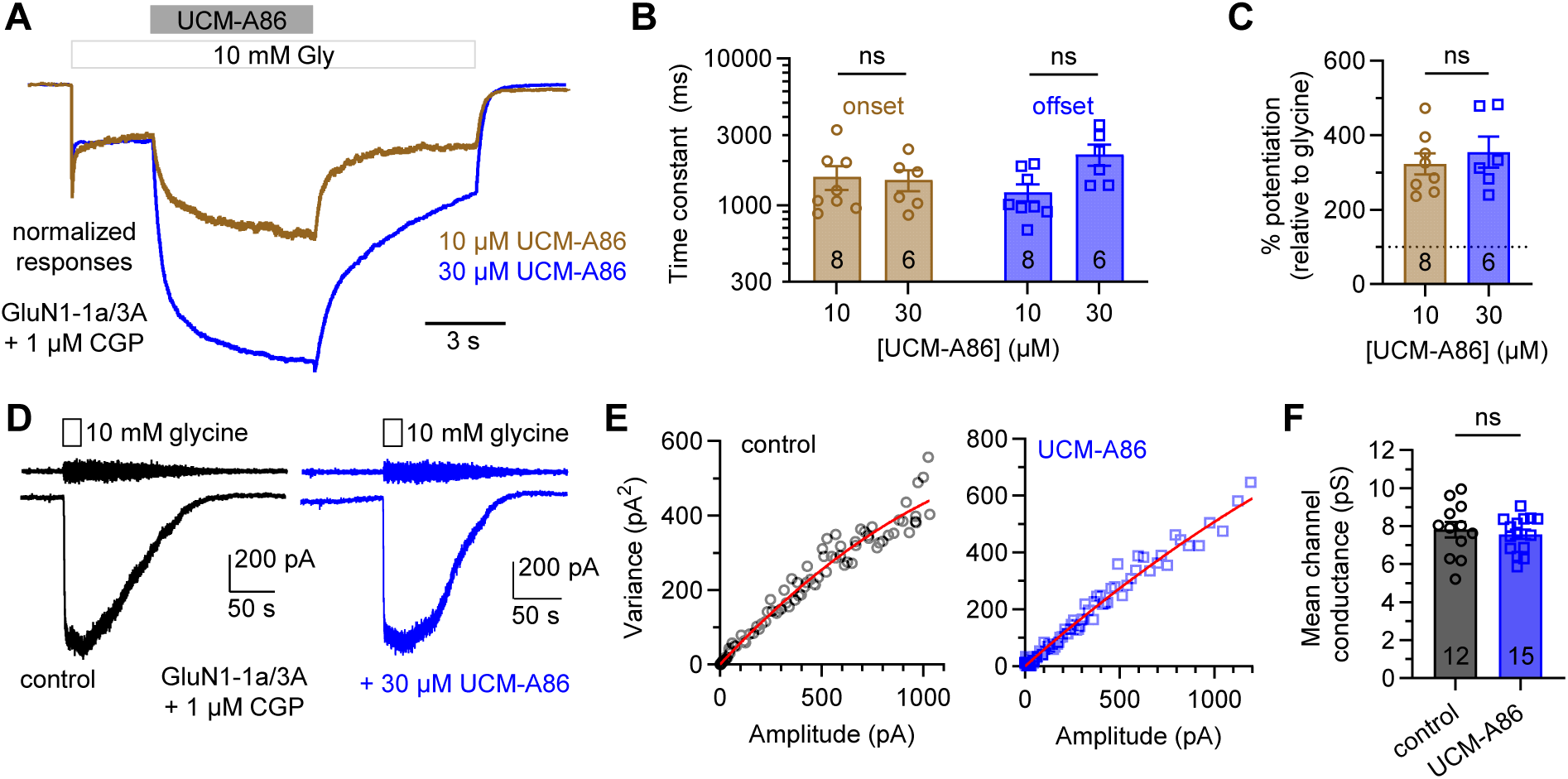
UCM-A86 modulation is mediated by a multistep mechanism with no effect on mean channel conductance. (A) Representative whole-cell patch-clamp recordings in the continuous presence of 1 µM CGP. Responses were activated by 10 mM glycine and modulated by co-application of either 10 µM or 30 µM UCM-A86. (B) Bar graph summarizing the weighted time constants for onset and offset of UCM-A86 potentiation. Significance determined by two-way ANOVA with Tukey post-test (ns, p > 0.05). (C) Bar graph summarizing potentiation relative to the response to glycine. Significance determined by unpaired t-test (ns, p > 0.05). (D) Representative whole-cell patch-clamp recordings of responses to bath-applied 10 mM glycine the continous presence of 1 µM CGP, but in the absence (vehicle, 0.15% DMSO) or presence of 30 µM UCM-A86. The same recording filtered at 1 Hz are shown above to illustrate noise arising from channel gating. (E) Current-variance plots from two different exposed to glycine alone (vehicle control) or glycine + UCM-A86. (F) Bar graph summarizing mean channel conductance measured from cells exposed to glycine alone (vehicle control) or glycine + UCM-A86. Significance determined by unpaired t-test (ns, p > 0.05).

**Table 3.**
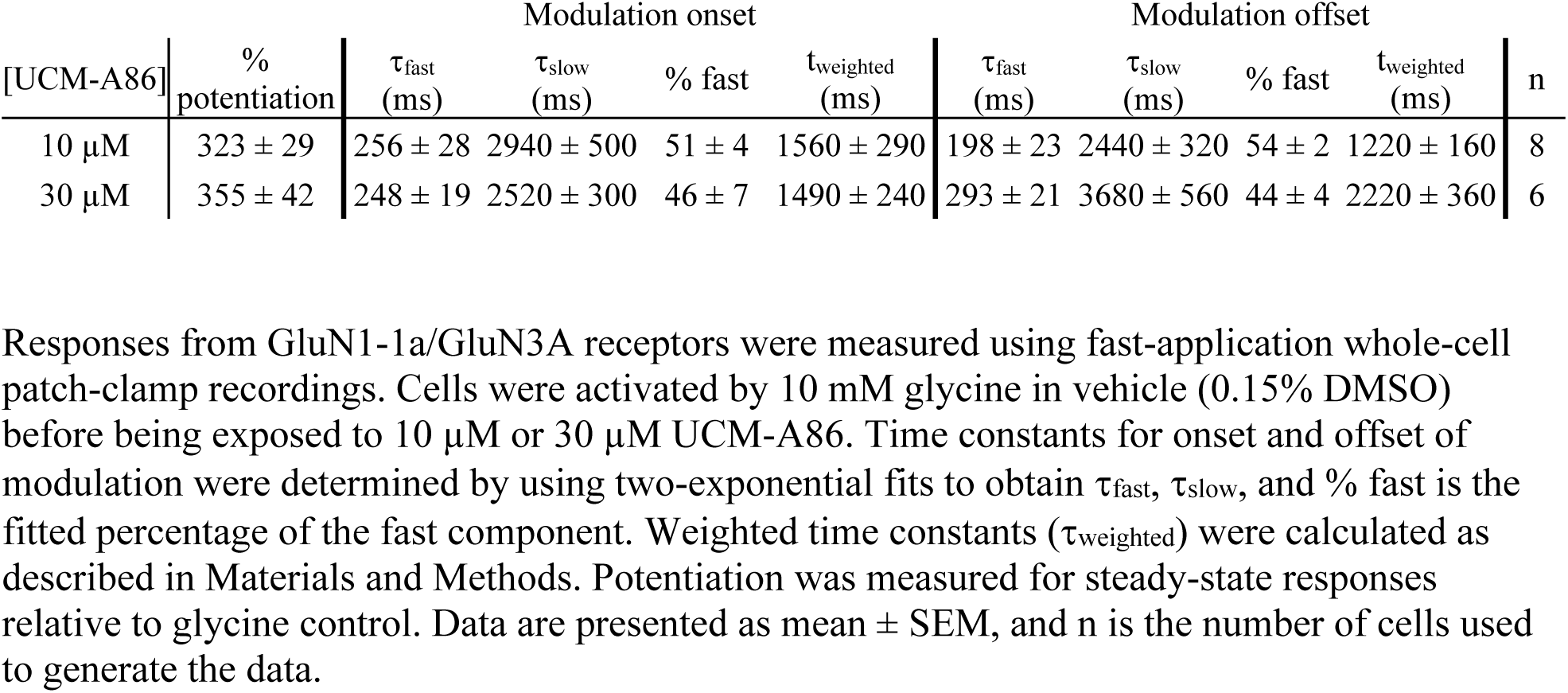
Kinetic parameters for onset and offset of modulation by UCM-A86.

| [UCM-A86] | %<br>potentiation | Modulation onset |  |  |  | Modulation offset |  |  |  | n |
| --- | --- | --- | --- | --- | --- | --- | --- | --- | --- | --- |
| | | $\tau_{\text{fast}}$<br>(ms) | $\tau_{\text{slow}}$<br>(ms) | % fast | $t_{\text{weighted}}$<br>(ms) | $\tau_{\text{fast}}$<br>(ms) | $\tau_{\text{slow}}$<br>(ms) | % fast | $t_{\text{weighted}}$<br>(ms) | |
| 10 $\mu\text{M}$ | 323 ± 29 | 256 ± 28 | 2940 ± 500 | 51 ± 4 | 1560 ± 290 | 198 ± 23 | 2440 ± 320 | 54 ± 2 | 1220 ± 160 | 8 |
| 30 $\mu\text{M}$ | 355 ± 42 | 248 ± 19 | 2520 ± 300 | 46 ± 7 | 1490 ± 240 | 293 ± 21 | 3680 ± 560 | 44 ± 4 | 2220 ± 360 | 6 |

Responses from GluN1-1a/GluN3A receptors were measured using fast-application whole-cell patch-clamp recordings. Cells were activated by 10 mM glycine in vehicle (0.15% DMSO) before being exposed to 10 $\mu\text{M}$ or 30 $\mu\text{M}$ UCM-A86. Time constants for onset and offset of modulation were determined by using two-exponential fits to obtain $\tau_{\text{fast}}$ , $\tau_{\text{slow}}$ , and % fast is the fitted percentage of the fast component. Weighted time constants ( $t_{\text{weighted}}$ ) were calculated as described in Materials and Methods. Potentiation was measured for steady-state responses relative to glycine control. Data are presented as mean ± SEM, and n is the number of cells used to generate the data.

### Mean channel conductance is not modulated by UCM-A86

We performed nonstationary noise analysis of responses from GluN1-1a/3A receptors activated by bath-application of 10 mM glycine in the continuous presence of 1 µM CGP using whole-cell patch-clamp electrophysiology (Fig. 7D-F). The mean channel conductance in bath-applied vehicle (0.15% DMSO) was 7.8 ± 0.4 pS (n = 12), which was not significantly different from the mean channel conductance of 7.6 ± 0.2 pS (n = 15) measured in the presence of 30 µM UCM-A86 (p > 0.05, unpaired t-test) (Fig. 7D-F). These results demonstrate that positive allosteric modulation by UCM-A86 is not accompanied by an increase in mean channel conductance of GluN1-1a/3A receptors.

## Discussion

Recently, negative allosteric modulators for GluN1/GluN3A receptors have been described (Zhu et al., 2020; Zeng et al., 2022), but positive allosteric modulators for GluN1/GluN3A NMDA receptors remain elusive. In this study, we characterized UCM-A86 as a PAM that selectively targets GluN3-containing over GluN2-containing NMDA receptors. Our findings establish UCM-A86 as the first selective GluN3 PAM, addressing a critical gap in available pharmacological tool compounds for studying these receptors. The detailed characterization of the macroscopic response time course in the presence and absence of UCM-A86 provides insights into the mechanism of action. A key finding of the study was that UCM-A86 modulation is facilitated by glycine binding, since UCM-A86 does not exert its full potentiating effects until after glycine binds the receptor. Experiments using pre-exposure of UCM-A86 before glycine activation suggest that robust UCM-A86 binding does not occur in the absence of glycine binding or channel gating. The dependency of UCM-A86 on glycine binding is consistent with further observations that UCM-A86 increases glycine potency, suggesting that UCM-A86 stabilizes the active glycine-bound conformation of the receptor.

The selectivity of UCM-A86 for GluN3-containing NMDA receptors facilitates investigations into the physiological roles of GluN3 subunits in neuronal function and provides a potential avenue for developing therapeutics that target GluN3-mediated processes without affecting the more abundant GluN1/GluN2 receptors. Our experiments in neuronal preparations further validate the selectivity and efficacy of UCM-A86. The significant potentiation of glycine-evoked currents in somatostatin-expressing interneurons of the somatosensory cortex, where GluN3A is highly expressed (Bossi et al., 2023), demonstrates that UCM-A86 can effectively modulate native GluN3-containing receptors. The absence of effects on AMPA or GluN1/2 NMDA receptor-mediated EPSCs in CA1 pyramidal neurons further substantiates the selectivity observed in heterologous systems.

The complex mechanism of action is demonstrated by kinetic measurements of onset and offset of UCM-A86 modulation, which reveal that the mechanism does not follow simple bimolecular interaction kinetics. This complexity could reflect multiple binding sites, cooperative interactions, or state-dependent accessibility of binding sites. Further structural and functional studies could elucidate the binding site of UCM-A86 on GluN1/GluN3 receptors and thereby provide additional insights into the precise mechanism of action. These studies would also enable structure-based design of UCM-A86 analogs with improved potency and efficacy, as well as potential selectivity between GluN3A and GluN3B subunits. Furthermore, additional studies may also reveal whether UCM-A86 can bind desensitized GluN1/GluN3 receptors in the absence of FATL mutations or CGP to prevent glycine binding to GluN1.

Our findings have potential implications for understanding the roles of GluN3 subunits in normal brain function and disease, including Huntington’s disease (Marco et al., 2013; Mahfooz et al., 2016), schizophrenia (Matsuno et al., 2015; Greenwood et al., 2016), addiction (Yuan et al., 2013; Christian et al., 2021), and fear- and anxiety-related behaviors (Conde-Dusman et al., 2021; Bossi et al., 2022; Pizzamiglio et al., 2025). The selective modulation of GluN3-containing receptors by UCM-A86 provides a new tool for investigating the role of these receptors in such pathologies. For instance, the developmental regulation of GluN3A expression suggests that selective modulation could be particularly relevant for neurodevelopmental disorders or for understanding critical periods in synaptic plasticity (Crawley et al., 2022). The identification of UCM-A86 as a GluN3-selective PAM opens new possibilities to modulate excitatory neurotransmission while avoiding the excitotoxicity associated with overactivation of calcium-permeable GluN1/GluN2 receptors, which is particularly relevant in conditions where GluN3A dysfunction has been implicated, such as schizophrenia or Huntington’s disease (Marco et al., 2013; Greenwood et al., 2016; Mahfooz et al., 2016).

In summary, given the role of GluN1/GluN3A receptors in normal and pathological brain function, there is growing interest in developing GluN3A subunit-specific allosteric modulators. The discovery and characterization of UCM-A86 as a selective PAM of GluN3-containing NMDA receptors represents a significant advance in our ability to study and potentially target these receptors. UCM-A86 therefore provides a valuable pharmacological tool for investigating GluN3 function and a novel starting point for the development of allosteric modulators targeting GluN3-containing NMDA receptors, which could have therapeutic potential in pathologies involving GluN3 dysfunction.

## Acknowledgements

We thank Gina C. Bullard for excellent technical assistance. We thank Atomwise Inc. (San Francisco, CA) for performing virtual screening and providing top hit compounds for functional evaluation at GluN1/GluN3A receptors through their academic collaboration program, Artificial Intelligence Molecular Screen (AIMS). The authors declare no conflicts of interest.

## Data Availability Statement

The authors declare that all the data supporting the findings of this study are available within the paper and its Supplemental Data.

## Authorship Contributions

*Participated in research design:* Benton, Johns, Diamant, Rouzbeh, Sørensen, Anderson, Secci, Zhang, Rau, Clausen, Hansen.

*Conducted experiments:* Benton, Johns, Diamant, Rouzbeh, Sørensen, Anderson, Secci, Zhang, Rau.

*Performed data analysis:* Benton, Johns, Diamant, Rouzbeh, Sørensen, Rau, Hansen.

*Wrote or contributed to the writing of the manuscript:* Benton, Johns, Diamant, Rouzbeh, Sørensen, Anderson, Secci, Zhang, Rau, Clausen, Hansen.

## Abbreviations

CGP: CGP 78608
CNS: central nervous system
NMDA: *N*-methyl-D-aspartate
PAM: positive allosteric modulator
UCM-A86: 6-(4-propan-2-ylphenyl)sulfanyl-3-(trifluoromethyl)-[1,2,4]triazolo[4,3-b]pyridazine.

## Footnotes

This work was supported by the National Institutes of Health [Grants P30GM140963, P20GM103474, R01NS097536, and R01NS116055]. No author has an actual or perceived conflict of interest with the contents of this article.

