## Supplemental Information for "UCM-A86 is a selective positive allosteric modulator of GluN1/GluN3 NMDA receptors"

**Supplemental material**

- Table S1 provides $\log EC_{50}$ values for potentiation of GluN1/3 receptors by UCM-A86.
- Supplemental Materials and Methods provide the synthesis route for UCM-A86.

**Supplemental Table S1. UCM-A86 modulation of GluN1/GluN3 receptor subtypes.**

| | GluN3A | | | | GluN3B | | | |
|---|---|---|---|---|---|---|---|---|
| | $LogEC_{50}$ (M) | Hill slope | % potentiation | n | $LogEC_{50}$ (M) | Hill slope | % potentiation | n |
| GluN1-1a$^{FATL}$ | -4.779 ± 0.064 | 1.6 | 396 ± 86 * | 12 | - | | | |
| GluN1-1b$^{FATL}$ | -4.893 ± 0.200 * | 1.6 | 367 ± 88 * | 13 | - | | | |
| GluN1-4a$^{FATL}$ | -4.928 ± 0.183 *,# | 1.7 | 392 ± 117 # | 16 | -4.689 ± 0.081 # | 1.6 | 195 ± 26 # | 11 |
| GluN1-1a$^{CGP}$ | -4.628 ± 0.122 | 1.7 | 558 ± 123 * | 13 | - | | | |
| GluN1-1b$^{CGP}$ | -4.686 ± 0.113 * | 1.6 | 511 ± 153 * | 10 | - | | | |
| GluN1-4a$^{CGP}$ | -4.669 ± 0.074 * | 1.7 | 436 ± 136 # | 11 | -4.719 ± 0.103 | 1.3 | 174 ± 11 # | 10 |

Concentration-response data for UCM-A86 at recombinant GluN1/3 NMDA receptor subtypes were measured using two-electrode voltage clamp electrophysiology. FATL indicates that desensitization was prevented by F484A+T518L mutations in the GluN1 subunit and CGP indicates that that desensitization was prevented by the continuous presence of 1 μM CGP-78608. Receptors were activated with 30 μM glycine in the absence and presence of increasing concentrations of UCM-A86. $EC_{50}$ and Hill slope values were determined by fitting the UCM-A86 concentration-response data to the Hill equation. Values are shown as mean ± SD. - indicates not determined, and n is the number of oocytes used as experimental sample size. Statistical analysis was performed by comparing all values using one-way ANOVA with Tukey posttest. * indicates significant difference between the approach to prevent desensitization (FATL or CGP) for the same GluN1 splice variant (e.g. GluN1-1a$^{FATL}$/3A versus GluN1-1a$^{CGP}$/3A) ($p < 0.05$). # indicates significant difference between corresponding values for GluN1-4a$^{FATL}$/3A versus GluN1-4a$^{FATL}$/3B or GluN1-4a$^{CGP}$/3A versus GluN1-4a$^{CGP}$/3B ($p < 0.05$).

**Synthesis of UCM-A86**

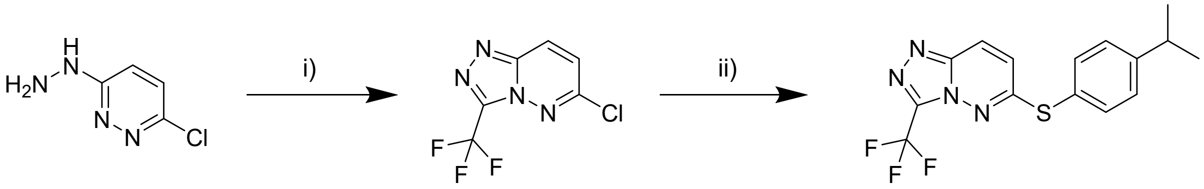

**Scheme 1.** Reagents and conditions: i) TFA, 110°C for 2 hours. ii) 4-isopropylbenzenethiol, $K_2CO_3$, DMF, RT for 2 hours.

**General procedures.**

Reactions in anhydrous conditions were carried out in flame-dried glassware under argon atmosphere. Chromatographic grade solvents were used, purified in Waters SG solvent purification system (only for DCM, THF, and DMF), while AcN, MeOH and EtOH were dried over molecular sieves (3 Å). Thin-layer chromatography was performed using TLC Silica gel 60 F254, Merck pre-coated plates. The plates were visualized under the UV lamp (254 and 365 nm) when needed stained with ninhydrin, $KMnO_4$, or 2,4-DNP. Flash chromatography was performed using silica gel 60 (40-64 µm) through a dry load method (celite).

NMR spectra were recorded on 400 or 600 MHz Bruker instruments. The FID files were analyzed with MestReNova, version 14.2.1. Signals are reported in ppm (δ) and the coupling constants are given in Hertz (Hz). Multiplet patterns are designated the following abbreviations, or combinations of these: m – multiplet, s – singlet, d – doublet, t – triplet, q – quartet, p – pentaplet, and h – sextet. Signal assignments were made from chemical shifts and indications were verified by HSQC (Heteronuclear Single Bond Correlation), COSY (Correlated Spectroscopy) and HMBC (Heteronuclear Multiple Bond Correlation) experiments.

Liquid Chromatography-Mass spectrometry (LC-MS) and the reported m/z values were obtained with an Agilent 6130 Mass Spectrometer instrument using electron spray ionization (ESI) coupled to an Agilent 1200 HPLC system (ESI-LCMS) with a $C^{18}$ reverse phase column (Zorbax Eclipse XBD-C18, 4.6 mm × 50 mm), autosampler and diode array detector, using a linear gradient of the binary solvent system of buffer A (milliQ $H_2O$:MeCN:formic acid, 95:5:0.1 v/v%) to buffer B (MeCN:formic acid, 100:0.1 v/v %) with a flow rate of 1 mL⁄min.

Analytical High-Performance Liquid-chromatography (HPLC) was carried out on an UltiMate HPLC system (Thermo Scientific) consisting of an LPG-3400A pump (1 mL/min), a WPS-3000SL autosampler, and a DAD-3000D diode array detector (210, 225, 254, 365 nm), using a Gemini-NX $C^{18}$ column (4.6 × 250 mm, 3 μm, 110 Å); Mobile phase A: $H_2O$ : TFA 100:0.1, v/v. Mobile phase B: MeCN : H2O : TFA 90:10:0.1, v/v/v. Data were acquired and processed using Chromeleon Software v. 6.80. Purity of the final compounds was assessed to be >95% using a gradient method 0-100% mobile phase B over 20 minutes.

**6-chloro-3-(trifluoromethyl)-[1,2,4]triazolo[4,3-b]pyridazine**

3-chloro-6-hydrazinopyridazine (2 g, 1 eq, 13.83 mmol) was added to a microwave vial and dissolved in 5.3 mL of TFA (5 eq, 69.17 mmol). The tube was sealed, and the mixture was stirred at 110°C for 2 hours. At reaction completion, the solvent was removed under reduced pressure. EtOAc (1.5 M) and potassium carbonate (95.6 g, 50 eq.) were added and an extraction in EtOAc/$H_2O$ was performed. The organic phase was dried over $MgSO_4$, filtered and evaporated to dryness to yield 6-chloro-3-(trifluoromethyl)-[1,2,4]triazolo[4,3-b]pyridazine as an off-white solid. $^1$H NMR (600 MHz, $CDCl_3$) δ 8.22 (d, J = 9.7 Hz, $^1$H), 7.33 (d, J = 9.7 Hz, $^1$H). $^{13}$C NMR (151 MHz, $CDCl_3$) δ 151.7, 144.5, 139.9 (q, J = 42.3 Hz), 126.6, 124.7, 118.1 (q, J = 271.1 Hz). Yield: 84.5%. m/z: 223 $[M+H]^+$

**6-((4-isopropylphenyl)thio)-3-(trifluoromethyl)-[1,2,4]triazolo[4,3-b]pyridazine (UCM-A86)**

6-chloro-3-(trifluoromethyl)-[1,2,4]triazolo[4,3-b]pyridazine (150 mg, 1 eq, 0.674 mmol), potassium carbonate (931 mg, 10 eq, 6.74 mmol) and 4-isopropylbenzenethiol (157 µL, 1.50 eq, 1 mmol) were added to a round bottom flask and dissolved in 1.3 mL of DMF. The mixture was stirred at room temperature for 4 hours. The mixture was dried, and an ether extraction was performed, the organic layer was washed with a saturated solution of $CaCl_2$, dried over $MgSO_4$. The solvent was evaporated in vacuo and the crude product purified by flash column chromatography (EtOAc/Heptane) to yield UCM-A86 as a yellowish powder. $^{1}H$ NMR (600 MHz, $CDCl_3$) δ 7.96 (d, J = 9.8 Hz, $^{1}H$), 7.54 (dd, J = 8.2, 1.6 Hz, $^{2}H$), 7.35 (dd, J = 8.2, 1.6 Hz, $^{2}H$), 6.98 (d, J = 9.8 Hz, $^{1}H$), 2.99 (hept, J = 7.0 Hz, $^{1}H$), 1.30 (d, J = 6.9 Hz, $^{6}H$). $^{13}C$ NMR (151 MHz, $CDCl_3$) δ 161.5, 152.3, 144.7, 139.5 (q, J = 41.8 Hz), 135.7, 128.3, 123.7, 123.3, 122.2, 118.2 (q, J = 270.8 Hz), 34.2, 23.9. Yield: 47%. m/z: 339 $[M+H]^+$

6-chloro-3-(trifluoromethyl)-[1,2,4]triazolo[4,3-b]pyridazine

$^{1}$H NMR (600 MHz, $CDCl_3$) δ 8.22 (d, $J$ = 9.7 Hz, 1H), 7.33 (d, $J$ = 9.7 Hz, 1H).

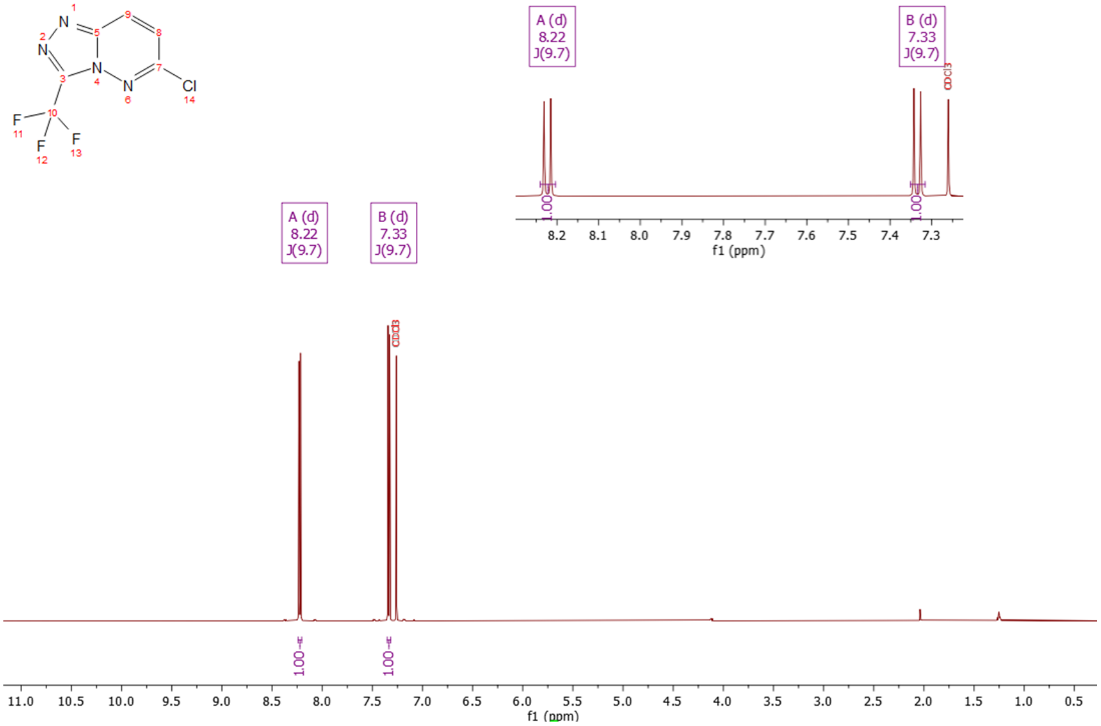


$^{13}$C NMR (151 MHz, $CDCl_3$) δ 151.7, 144.5, 139.9 (q, $J$ = 42.3 Hz), 126.6, 124.7, 118.1 (q, $J$ = 271.1 Hz).

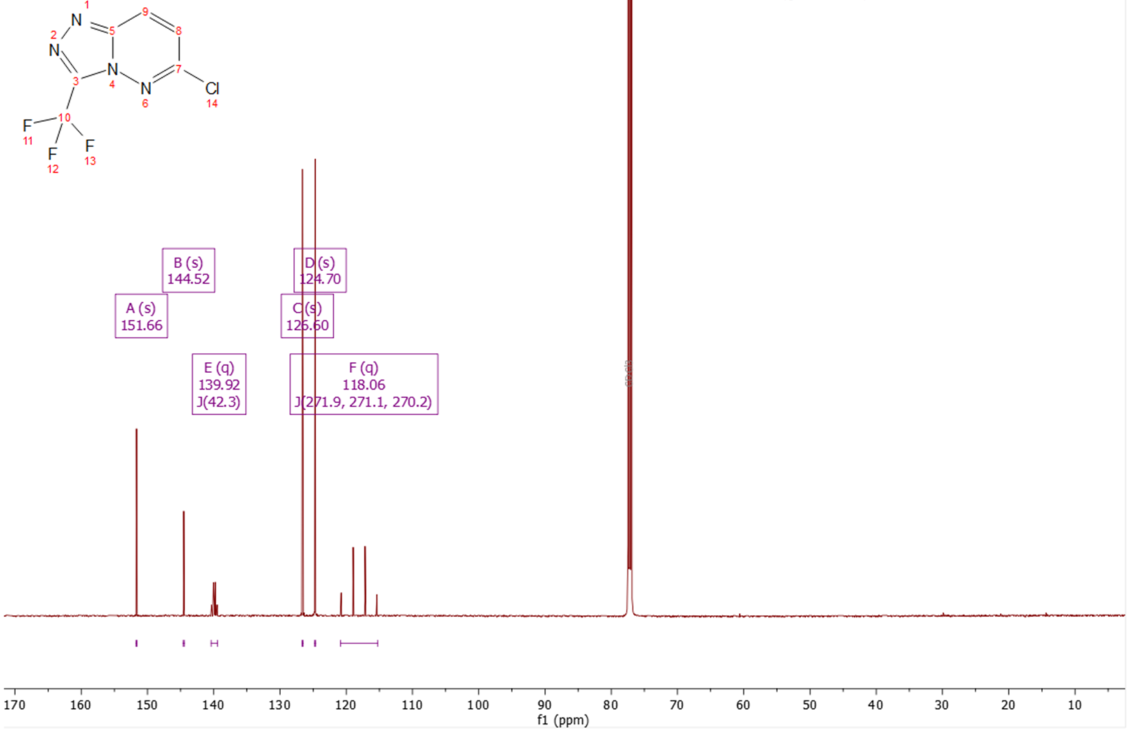

6-((4-isopropylphenyl)thio)-3-(trifluoromethyl)-[1,2,4]triazolo[4,3-b]pyridazine (UCM-A86)

$^{1}H$ NMR (600 MHz, $CDCl_3$) δ 7.96 (d, $J$ = 9.8 Hz, 1H), 7.54 (dd, $J$ = 8.2, 1.6 Hz, 2H), 7.35 (dd, $J$ = 8.2, 1.6 Hz, 2H), 6.98 (d, $J$ = 9.8 Hz, 1H), 2.99 (hept, $J$ = 7.0 Hz, 1H), 1.30 (d, $J$ = 6.9 Hz, 6H).

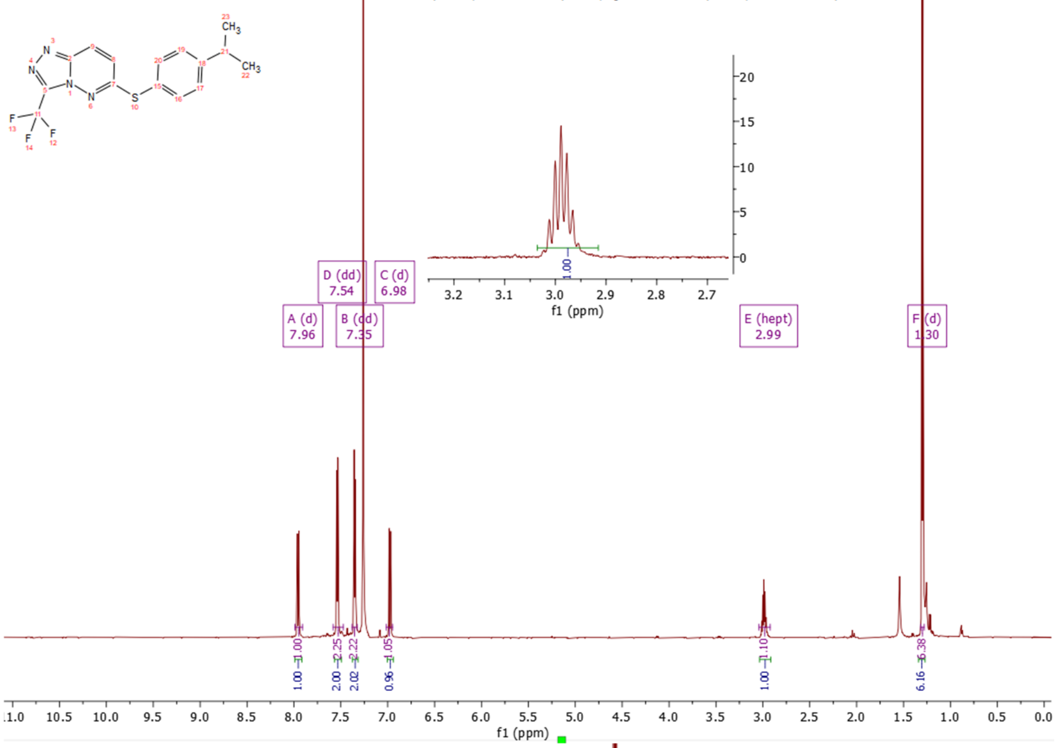


$^{13}C$ NMR (151 MHz, $CDCl_3$) δ 161.51, 152.26, 144.68, 139.48 (q, $J$ = 41.8 Hz), 135.66, 128.29, 123.73, 123.27, 122.19, 118.19 (q, $J$ = 270.8 Hz), 34.24, 23.93.

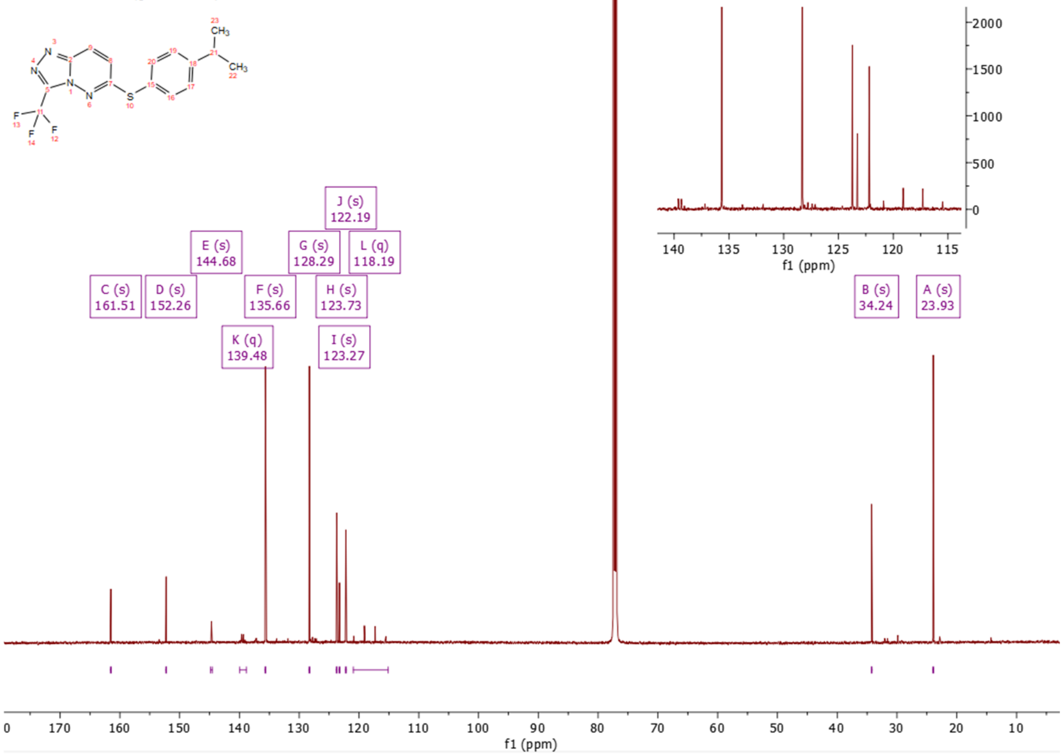
